# An AINTEGUMENTA phospho-switch controls bilateral stem cell activity during secondary growth

**DOI:** 10.1101/2024.06.20.599823

**Authors:** Wei Xiao, Ling Yang, Hengqi Ji, David Molina, Houming Chen, Shan Yu, Yingjing Miao, Dagmar Ripper, Shulin Deng, Martin Bayer, Bert De Rybel, Laura Ragni

## Abstract

Plant stem cells have the remarkable ability to give rise to distinct tissues and organs throughout development. Two concentric cylinders of actively dividing stem cells are the main drivers of radial thickening during secondary growth, each producing distinct vascular cell types towards the inside and the outside. While the molecular mechanisms underlying their initiation have been well studied, it remains unclear how these stem cell layers determine which cell type is generated. Here, we demonstrate that the cross-talk between the ERECTA (ER) receptor pathway and auxin signalling controls the amount and choice of output tissue in these two stem cell regions. Mechanistically, we show that the ER pathway phosphorylates the transcription factor AINTEGUMENTA, thereby modulating bilateral stem cell activity and favouring the differentiation of only one type of cell. Our results thus show that the phosphorylation status of ANT regulates root girth and biomass accumulation by influencing bilateral stem cell production. This highlights the critical role of post-translational modifications in plant growth and development.

**Significance:** This study uncovers a critical regulatory mechanism governing bilateral stem cell activity during plant radial growth. Stem cells in the cambia proliferate and differentiate into specialized tissues like wood and cork, key sites of biomass accumulation. By exploring the interplay between the ERECTA (ER) receptor and auxin signalling, we reveal how these pathways modulate stem cell differentiation and radial thickening through the phosphorylation of the transcription factor AINTEGUMENTA (ANT). Phosphorylation status of ANT modulates stem cell activity, dictating the production of specific vascular cell types like wood thus controlling root girth. This research underscores the importance of post-translational modifications in plant development and provides a novel regulation underpinning stem cell differentiation, with significant implications for enhancing biomass accumulation as wood.

## Introduction

During secondary growth, plant organs undergo a tremendous increase in girth. This is required to provide sufficient mechanical strength during growth and simultaneously increases the capacity of the vascular tissues. In woody plants, most biomass is produced by organs undergoing secondary growth in the form of wood and cork^1,2^. The growth process of secondary growth primarily relies on proliferation within two lateral stem cell zones called the vascular cambium and cork cambium^3–5^. In the Arabidopsis root, these stem cells are organized as two concentric cylinders of dividing cells: the vascular cambium forms the xylem toward the interior and phloem towards the exterior, and all together are referred as the secondary vascular tissues, while the cork cambium produces the periderm with the parenchymatic phelloderm towards the inside and the cork (phellem) towards the outside (**Fig S1A**)^6–8^. Despite their structural independence, the relative amount of secondary vascular tissues and periderm produced toward the inside or outside varies in response to plant secondary growth or environmental cues^9,10^. However, the molecular mechanisms that control bilateral output production remain elusive.

It is, however, known that phytohormones regulate the self-renewing and differentiation of stem cells in plants^11–13^. For example, recent studies have shown that auxin maxima in proximity to the xylem vessels are required for the initiation and maintenance of vascular cambium activity^14^. Downstream of auxin, AUXIN RESPONSE FACTOR 5/ MONOPTEROS (ARF5/MP) activates the expression of *WUSCHEL-RELATED HOMEOBOX 4* (*WOX4*) and *CLASS III HOMEODOMAIN-LEUCINE ZIPPER* (*HD-ZIPIII*) genes, which regulate cell proliferation^14,15^. During periderm development, auxin accumulates in the cork cambium and controls its initiation by regulating the expression of *WOX4* and *BREVIPEDICELLUS* (*BP*)^7^. Furthermore, activation of *BP* and *WOX4* significantly enhances cambial proliferation and leads to an increase in the number of periderm layers^7,15^. Cytokinins (CK) also regulate secondary growth by promoting cell proliferation^16^ via transcriptional regulators such as *LATERAL ORGAN BOUNDARIES DOMAIN 3* (*LBD3*), *LBD4*^17^, and *AINTEGUMENTA* (*ANT*)^18^. The member of the AP2 transcription factor (TF) family, *ANT,* is first described to control flower organ initiation and maintenance of meristematic competence of plants^19–21^. Likewise, it works with cell-cycle player CYCD3;1 to promote cell divisions in the root vascular cambium^18^. Besides phytohormones, several peptides and their respective receptor kinases act as signalling hubs to modulate secondary growth. For example, the leucine-rich receptor-like kinases (LRR-RLK) of the ERECTA (ER) and PHLOEM INTERCALATED WITH XYLEM/TDIF RECEPTOR (PXY/TDR) families control procambium cell divisions^7,14,15,22–30^. Moreover, in hypocotyls, ER recognizes its ligands EPIDERMAL PATTERNING FACTOR (EPF)/EPFL-LIKE (EPFL4/6) and functions in xylem initiation^24^.

In this study, we show that auxin signalling triggers ER-dependent phosphorylation of ANT. The phosphorylation status of ANT defines whether more cells are generated toward the inside or the outside of the vascular and cork cambium. This ANT-dependent phospho-switch thus serves as a mechanism to precisely control which conductive and protective tissue types are generated by the lateral stem cell niches during secondary growth.

## Results

### Auxin signalling controls the type of secondary tissues produced by lateral stem cell regions

To gain insights into the proliferation dynamics of the vascular and cork cambium stem cell regions involved in secondary growth, we initially performed a quantitative description of these stem cell regions during development (**Fig 1A-B** and **S1A-B**). We first focused our attention on the vascular cambium stem cell region by quantifying the number of xylem cells (representing the output toward the inside of the vascular cambium) and the total number of secondary vascular tissue cells (representing the combined bilateral output of the vascular cambium). We sampled the uppermost parts of roots (0.5 cm from hypocotyl) and quantified the aforementioned features before (6-day-old roots), during (8-day-old roots), and after (12-day-old roots) the onset of secondary growth. As an additional control, we included 20-day-old roots with prominent secondary growth (**Fig 1A**). The number of secondary vascular cells and secondary xylem cells steadily and significantly increased during secondary growth (**Fig 1A-B**), which led to a clear expansion of the root girth. The number of secondary xylem cells most prominently increased after day 12, confirming previous results in the hypocotyl region^31^. Although a 2-day treatment with the synthetic auxin 1-naphthaleneacetic acid (NAA) significantly increased the total number of secondary vascular tissue cells, there was no significant effect on the number of secondary xylem cells^7,14,15^ (**Fig 1C-D**). These results indicate that auxin treatment promotes vasculature tissue proliferation but not secondary xylem differentiation.

**Fig. 1.**
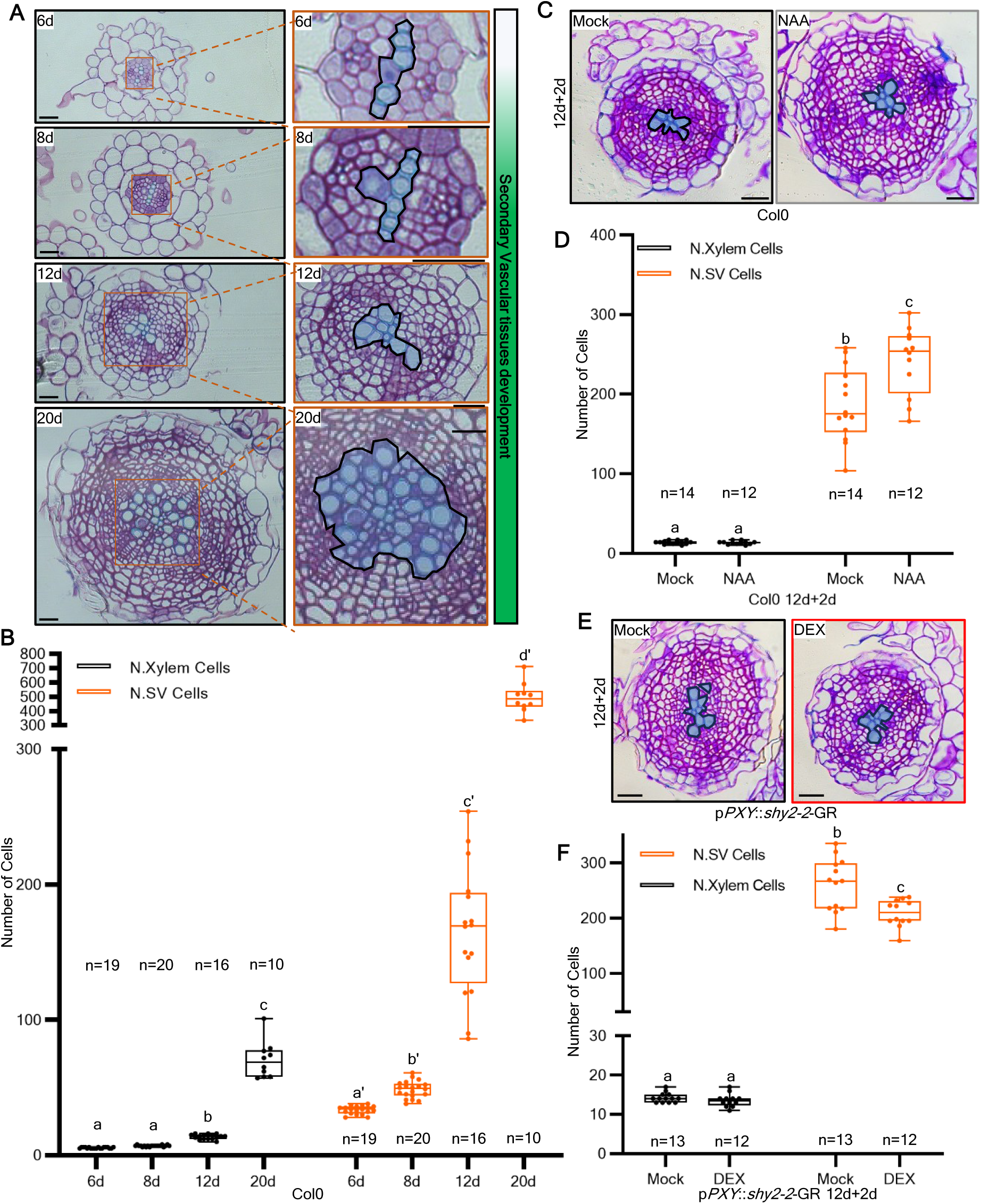
Auxin signalling preferential controls the specific output of vascular cambium. **(A)**, Secondary growth progression in the Arabidopsis root. Cross-sections of the most mature part of 6-day, 8-day, 12-day, and 20-day roots (*in vitro*). In the left panels, the orange squares indicate the magnification area shown in the right panels. Xylem cells are highlighted in blue. **(B)**, Quantification of secondary vascular (SV) tissue and xylem cell numbers of the experiment shown in (Fig1A). **(C)**, Cross-sections (plastic embedding) of the uppermost part of Col0 roots. 12-day-old plants were transferred to either Mock (left panel) or 1μM NAA (right panel) plates for 2 days. **(D)**, Quantification of the number of secondary vascular tissues and xylem cells in (Fig1C). **(E)**, Cross-sections (plastic embedding) quantification of the uppermost part of p*PXY*::*shy2-2*-GR roots (p*MYB84*::NLS-3xGFP and W131Y background); 12-day-old plants were transferred to either Mock (left panel) or 10 μM DEX (right panel) plates for 2 days. **(F)**, Quantification of the number of secondary vascular tissues and xylem cells in (Fig1E). The xylem vessels are highlighted in blue. Black scale bars: 20μm. Count data was modelled with a generalized mixed model. Here the Poisson mix model or Poisson model was selected. P-values were adjusted using Holm-Bonferroni correction.

To consolidate these results, we next specifically impaired canonical auxin signalling in the vascular cambium, in an inducible manner using a p*PXY*::*shy2-2*-GR construct^7,32^. Repression of auxin signalling in the vascular cambium suppressed the number of secondary vascular tissues (**Fig 1E-F**). To specifically dissect which cell types were affected, we quantified the number of xylem vessels (xylem cells are identified by prominent secondary cell wall accumulation), the number of vascular cambium cells (by expression of the p*PXY*::er-CFP marker), and phloem cells (by subtraction). We observed that impairing auxin signalling in the vascular cambium had a negative effect on the number of phloem and vascular cambium cells in secondary vascular tissues without a significant effect on the number of xylem cells (**Fig S1C**). To investigate whether a similar response could be observed in the cork cambium stem cell region, we next quantified the number of cork cells, the total number of periderm cells, and the number of periderm layers^7^ during (12-day-old roots) and after (20-day-old roots) periderm formation (**Fig S1D-F**). Although the number of periderm layers increased over time, the number of cork layers remained constant, with only a single cork layer (**Fig S1D**). Exogenous auxin treatment significantly stimulated cork cambium cell proliferation but exhibited no discernible effect on cork cell formation (**Fig S2A-D**). Next, we specifically impaired canonical auxin signalling in the cork cambium in an inducible manner using a p*PER15*::*slr*-GR construct^7,32^. Blocking auxin signalling overall resulted in fewer periderm layers, but an increase in the number of cork cells (**Fig S2E-H**). Taken together, these results suggest that auxin is not only required for cell proliferation in the vascular and cork cambium^7,14^ but also influences the relative amounts of secondary tissues that are derived from both stem cell regions.

### ERECTA controls directional vascular cambium activity downstream of auxin signalling

To uncover how auxin signalling might influence vascular and cork cambium activities on a molecular level, we performed bulk RNA sequencing (RNA-seq) on 12-day-old roots of the p*PXY*::*shy2-2*-GR and p*PER15*::*slr*-GR lines grown on mock or DEX (**Fig S3A**). Among the subset of genes significantly repressed by DEX treatment in both lines, we found several known regulators, including *WOX4*, *BP*, *ER, LBD3*, *PIN4* and *ANT* (**Fig S3B-C**). These have been demonstrated to promote cork and or vascular cambium activity^7,14,15,18,33^. In Arabidopsis, the ER receptor kinase family comprises three genes: *ER, ER-LIKE1* (*ERL*1) and *ERL2*. They play a role in many aspects of plant development, including maintaining vascular cambium activity^27,29,30^. However, little is known about the underlying mechanism of ER-dependent signalling in this developmental context. As shown for the hypocotyl and stem single*, er, erl1* and *erl2* mutants do not show any vascular phenotypes^24,27,34^. Furthermore, auxin responses were comparable to WT, probably due to compensation mechanisms inside the family (**Fig S3D-E**)^24^. To reduce functional redundancy within ER signalling, we next analysed the *er erl2* double mutant. Our quantitative analysis revealed that *er erl2* exhibited a trend toward increased xylem vessel number (**Fig 2A-B**). Additionally, *er erl2* double mutant did not respond to auxin compared to a WT control (**Fig 2A-B**), suggesting a potential connection between the ER pathway and auxin signalling. To further explore the functions of ER signalling during secondary growth, we investigated the effect of ER/ERL signalling inactivation in the cambia. However, the *er erl1 erl2* triple mutant is severely affected in overall growth^29^, preventing the analysis of secondary growth and hampering the identification of direct secondary growth phenotypes. To overcome these limitations, we employed tissue-specific expression of an anti-GFP nanobody, which targets the ER-YPet fusion protein for degradation. We introduced this construct in an *er erl1 erl2* triple mutant complemented with a p*ER*::ER-YPet construct^35^. Reducing expression levels of ER-YPet in the vascular cambium using this genetic tool led to a reduction in secondary vascular tissues without changing the number of secondary xylem cells (**Fig S3F-G**).

**Fig. 2.**
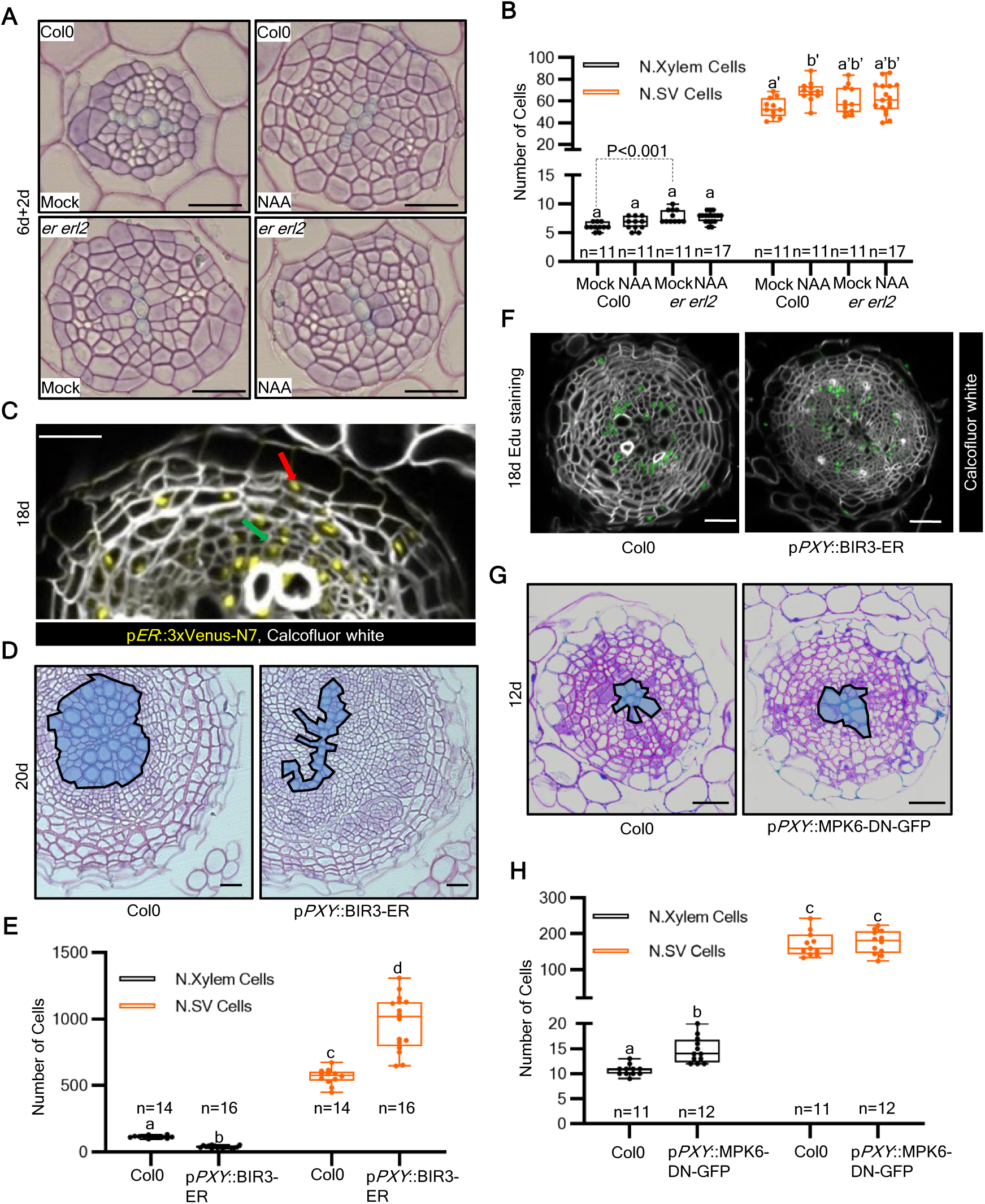
The ERECTA and MPK6 kinase trigger the directional vascular cambium activity. **(A)**, Cross-sections (plastic embedding) of the uppermost part of Col0 and *er erl2* roots. 6-day-old plants were transferred for 2 days on Mock (left panels) or 1μM NAA (right panels) plates. **(B)**, Quantification of the number of secondary vascular (SV) tissues and xylem cells in (Fig2A). **(C)**, Vibratome-sections of the uppermost part of 18-day-old p*ER::*3xVenus-N7 root. The red arrow indicates the cork cambium and the green arrow indicates the vascular cambium. **(D)**, Cross-sections (plastic embedding) of the uppermost part of 20-day-old Col0 (left panel) and p*PXY*::BIR3-ER (right panel) roots. The xylem vessels are highlighted in blue. **(E)**, Quantification of the number of secondary vascular tissues and xylem cells in (Fig2D). **(F)**, Edu staining of vibratome-sections of the uppermost part of 20-day-old Col0 (left panel) and p*PXY*::BIR3-ER (right panel) roots. **(G)**, Cross-sections (plastic embedding) of the uppermost part of 20-day-old Col0 (up-left panel) and p*PXY*::MPK6-DN-GFP (up-right panel) roots. The xylem vessels are highlighted in blue. **(H)**, Quantification of the number of secondary vascular tissues and xylem cells in (Fig2G). Black and white scale bars: 20μm. Count data was modelled with a generalized mixed model. Here the Poisson mix model was selected. P values were adjusted using Holm-Bonferroni correction. Count data was also analysed using the Student’s T-test.

Given that our transcriptomic data and qPCR results suggested that auxin promotes the expression of *ER* but not *ERL1* and *ERL2* (**Fig S3B-C and S4A**), we further focused our study on *ER*. Thus, we examined *ER* expression pattern (p*ER*::NLS-3xVenus) in roots undergoing secondary growth (**Fig S4B**)^7^. *ER* expression at the onset of secondary growth was found in the forming vascular cambium and pericycle cells (**Fig S4B**), while at later stages in both vascular and cork cambium (**Fig 2C**), which resembles the auxin signalling pattern^7,14^. To further dissect a putative role of ER in specifying the output of the vascular cambium, we took advantage of a previously published BAK1-INTERACTING RECEPTOR-LIKE KINASE3 (BIR3)-ER chimera protein^36^. This line allows the activation of the ER pathway in a ligand-independent way. Driving the chimeric BIR3-ER protein from a vascular cambium specific promoter (p*PXY*::BIR3-ER) triggered cell proliferation in the vascular cambium while the number of secondary xylem cells was repressed (**Fig 2D-E** and **S4C**). Given that the xylem defects might be due to the fact that *PXY* expression in the vasculature cambium is biased toward the proximal (xylem) side of the cambium and encompasses differentiating xylem, we drove the expression of *BIR3-ER* from a promoter that is expressed in the distal domain of the vascular cambium and in early phloem and overlaps with p*PXY* only in the stem cells of the vascular cambium (*SUPPRESSOR OF MAX2 1-LIKE 5*/*SMXL5*)^37,38^. p*SMXL5*::BIR3-ER roots showed similar phenotypes to p*PXY*::BIR3-ER roots in terms of xylem cell differentiation and secondary vascular tissue formation (**Fig S4D).** In addition, in both p*PXY*::BIR3-ER and p*PXY*::BIR3-ER x p*PXY*::er-CFP lines, we observed a broader distribution of dividing vascular cambium cells in secondary vascular tissues as well as ectopic secondary xylem cells (**Fig 2F** and **S4E-F**), suggesting that these phenotypes are triggered by the activation of ER signalling in vascular cambium. Collectively, our findings indicate that activation of ER signalling preferentially activates cell proliferation and directs the production of the vascular cambium toward the outside of the root, resulting in more secondary vascular cambium cells.

### ER phosphorylates ANT via the MPK6 kinase to determine vascular cambium activity

To explore the potential cross-talk between auxin signalling and ER pathway, we tested whether ER activates well-known auxin-responsive cambial regulators (**Fig S3B-C** and **S5A**)^7,15^. *ANT* expression was detected in pericycle, cork cambium and vascular cambium cells (**Fig 3A** and **S5B**)^15^ and *ANT* levels were specifically increased in p*PXY*::BIR3-ER (**Fig S5A**) roots, suggesting that ANT might be activated by the ER pathway.

**Fig. 3.**
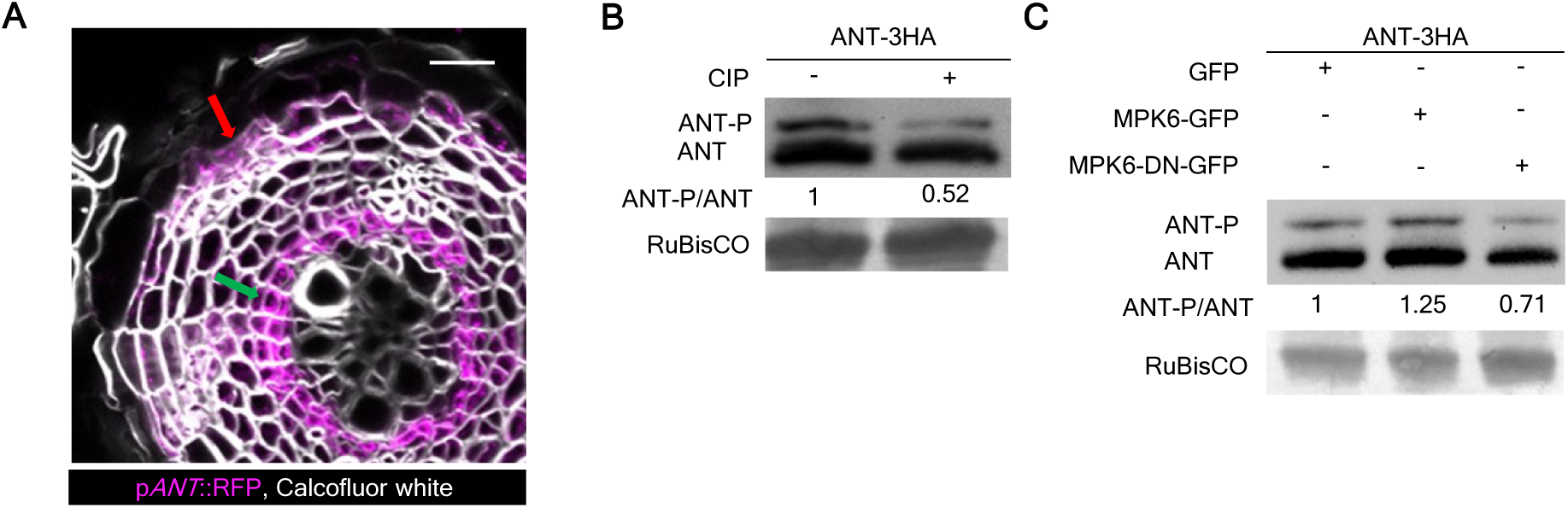
ER phosphorylates ANT via the MPK6 kinase. **(A)**, Vibratome-sections of the uppermost part of 18-day-old p*ANT*::RFP roots. The red arrow indicates the cork cambium and the green arrow indicates the vascular cambium. **(B)**, *In vivo* phosphorylation assays in *N. benthamiana.* Protein extracts, with or without treatment with Calf-intestinal alkaline (CIP), were analysed on a Phos-tag SDS-PAGE gel and probed with an anti-HA antibody. **(C)**, *In vivo* phosphorylation assays in *N. benthamiana.* p*35S*::ANT-3xHA were transiently expressed in leaves together with either p*35S*::MPK6-GFP or p*35S*::MPK6-DN-GFP (its dominant negative version). Protein extracts were analysed on a Phos-tag SDS-PAGE gel and probed with an anti-HA antibody. The relative concentration of phosphorylated ANT protein and unphosphorylated ANT protein was calculated using Fiji. White scale bars: 20μm.

Given that the ER pathway acts by phosphorylating downstream TF targets^39,40^, we next explored whether this could occur via the conserved YODA-MKK4/5-MPK3/6 MAPK cascade. This MAP kinase cascade has been shown to function as an integral part of the ER signalling pathway in other developmental contexts, such as embryogenesis and stomatal development^29,30,35,41–44^. Moreover, *MPK6* expression pattern overlaps with *ER* and *ANT* in the cork and vascular cambium (**Fig S5C**). Thus, to understand whether MPK6 acts as a downstream component of ER in regulating root secondary growth, we generated lines expressing a dominant negative version of MPK6^45^ in the vascular cambium (p*PXY*::MPK6-DN-GFP). Roots harbouring this construct showed a significant increase in the number of secondary xylem cells without impacting overall secondary vascular tissues (**Fig 2G-H**). As inactivating the MPK6 pathway resembles *er/erl* deficiency and results in the opposite effect compared to constitutively activating ER (p*PXY*::BIR3-ER) (**Fig 2A-B and D-E**), our results suggest that the ER pathway might activate MPK6 to favour directional vascular cambium activity, limiting secondary xylem cells formation.

To next explore whether ER might phosphorylate ANT via MPK6, we first conducted an *in silico* analysis based on the putative MPK6 target motif to identify potential phosphorylation sites in ANT (**Fig S5D**)^46–48^. Four putative phospho-sites were predicted in ANT (S113, S202, T496 and S526) (**Fig S5D**). To assess ANT phosphorylation *in vivo*, we then transiently expressed HA-tagged ANT in *Nicotiana benthamiana* leaves in the presence and absence of the Calf-intestinal alkaline phosphatase (CIP) and found that the phosphorylated form of ANT was reduced after CIP treatment (**Fig 3B**), indicating that ANT is phosphorylated in *N. benthamiana* leaves. Phosphorylation of ANT was increased in the presence of MPK6 and reduced when co-expressed with a dominant negative MPK6 version (**Fig 3C**), suggesting that MPK6 can phosphorylate ANT *in planta*. To further support these results, we expressed in *N. benthamiana* leaves, a phospho-dead version of ANT, all putative phospho-sites were replaced by alanine (S113A, S202A, T496A and S526A, called ANT-4A) and a phospho-mimic version (ANT-4D) with aspartic acid substitutions at all four sites (S113D, S202D, T496D and S526D). In a Phos-tag gel, we observed only one band for ANT-4A protein corresponding to the non-phosphorylated version, whereas ANT-4D protein mimicked the phosphorylated version (**Fig S5D**-**E**)^49^. Taken together, our data suggest that ER phosphorylates ANT via the MPK6 kinase pathway.

To next determine if ANT plays a role in modulating vascular cambium activity, we studied the effect of specifically inducing ANT in the vascular cambium (p*PXY*::ANT-GR on DEX). *ANT* induction inhibited vascular cambium differentiation toward the xylem, resulting in fewer secondary xylem cells (**Fig 4A-B** and **S6A-B**), highlighting a role for ANT in regulating directional vascular cambium output. Then, we studied the effect of ANT phosphorylation status by comparing the effect of ANT and the ANT-4A expression in the vascular cambium (p*PXY*::ANT-GFP or p*PXY*::ANT-4A-GFP). Constitutive overexpression of *ANT* in the vascular cambium resulted in fewer xylem vessel cells (**Fig S6C-D**), while expression of the phospho-dead version resulted in a milder phenotype (**Fig S6C-D**), highlighting a role for ANT phosphorylation in regulating directional vascular cambium activity. To further understand the functions of ANT, we generated an inducible chimeric repressor construct by fusing ANT with an EAR-repression domain (SRDX) (p*PXY*::ANT-SRDX-GR)^50^. Induction of *ANT-SRDX* (p*PXY*::ANT-SRDX-GR on DEX) in the vascular cambium triggered a reduction of secondary vascular tissues while xylem cell number was increased (**Fig 4C-D** and **S6E-F**), confirming that ANT balances vascular cambium output.

**Fig. 4.**
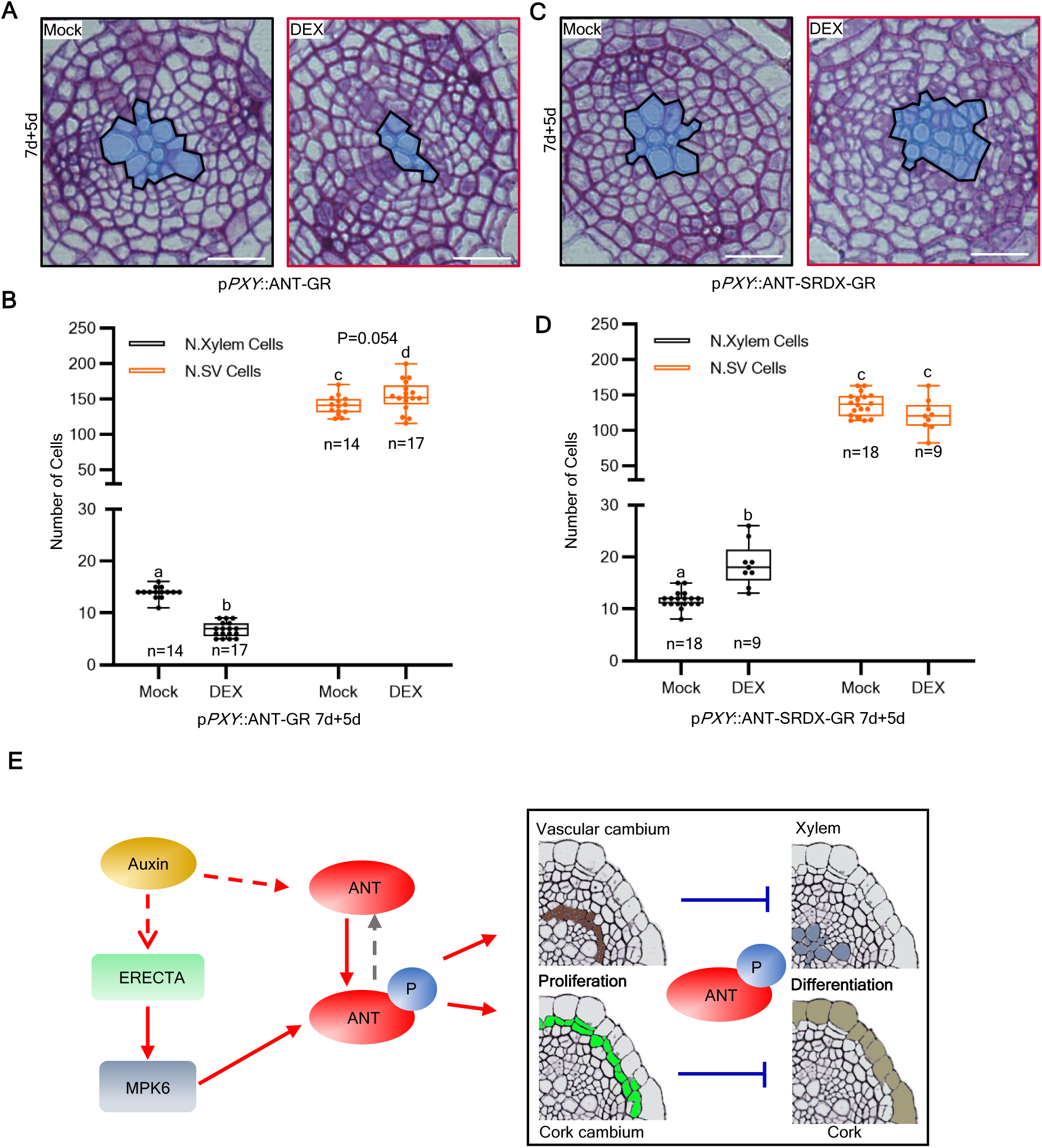
ANT determines vascular cambium activity by repressing xylem cell formation. **(A)**, Cross-sections (plastic embedding) of the uppermost part of 12-day-old p*PXY*::ANT-GR roots; 7-day-old plants were transferred to either Mock (left panel) or 10 μM DEX (right panel) plates for 5 days. **(B)**, Quantification of the number of secondary vascular (SV) tissues and xylem cells shown in (Fig4A). **(C)**, Cross-sections (plastic embedding) of the uppermost part of 12-d-old p*PXY*::ANT-SRDX-GR roots; 7-day-old plants were transferred to either Mock (left panel) or 10 μM DEX (right panel) plates for 5 days. **(D)**, Quantification of the number of secondary vascular tissues and xylem cells shown in (Fig4C). **(E)**, Model explaining cambia output specificity. White scale bars: 20μm. Count data was modelled with a generalized mixed model. Here the Poisson mix model was selected. P values were adjusted using Holm-Bonferroni correction. Count data was also analysed using the Student’s T-test.

To corroborate our hypothesis that ANT is phosphorylated by the ERECTA pathway, we tested whether the effect of ER constitutive activation could be suppressed by reducing ANT function. Induction of p*PXY*::ANT-SRDX-GR in p*PXY*::BIR3-ER lines was able to rescue the xylem deficiency phenotype and secondary vasculature tissue overproduction of p*PXY*::BIR3-ER (**Fig S3C** and **S7A**), indicating that ANT is downstream of ER signalling. Additionally, we studied the effect of inducing *ANT*, *ANT* phospho-dead (*ANT-4A*), and ANT phospho-mimic (*ANT-4D*) in a background with reduced ER signalling (p*PXY*::NSlmb-vhhGFP4 in the *er erl1 erl2* triple mutant complemented with p*ER*::ER-YPet mutants). Only the induction of *ANT-4D* inhibited xylem formation in ER limiting conditions, while the induction of *ANT* and *ANT-4A* did not alter the number of xylem vessels and vascular tissues (**Fig S7B-E**), further supporting that ANT phosphorylation is mediated by ER signalling and it is needed for ANT function. Together, these results suggest that ER-MPK6-dependent phosphorylation of ANT controls directional vascular cambium activity and that ANT is an important factor controlling secondary xylem formation.

### The ANT-dependent phospho-switch acts in both lateral cambia

Our results showed that both *ANT* and *ER* are expressed in the cork cambium and in the pericycle (the tissue that by divisions, will give rise to the cork cambium) (**Fig 2C**, **Fig 3A**, **Fig S4B**, and **S5D**). Furthermore, the expression was downregulated by blocking auxin signalling in the cork cambium (**Fig S3B-C**). Therefore, we hypothesized that a similar ER-ANT-dependent phospho-switch might be active in the periderm to control the directional output of the cork cambium. Indeed, activation of ER specifically in the periderm (p*PER15*::BIR3-ER) increased the number of the periderm cells and layers, while it showed no impact on the number of cork layers (**Fig S8A-D**). Reducing expression levels of ER, specifically in the cork cambium using an anti-GFP nanobody approach (p*PER15*::NSlmb-vhhGFP4 in the *er erl1 erl2* triple mutant complemented with p*ER*::ER-YPet) decreased the number of periderm layers and number of periderm cells without affecting the number of cork layers or cork cells (**Fig S8E-H**). Moreover, induction of *ANT,* specifically in the cork cambium (p*PER15*::ANT-GR on DEX), resulted in roots with an increased number of periderm layers and cells without alteration of cork layer and cell numbers (**Fig S9A-D**). These results thus suggest a similar role for ER in regulating ANT to control directional cork cambium activity. In summary, we proposed that both the vascular and cork cambium share a core regulatory mechanism based on an ER-dependent phospho-switch of ANT to control directional cambium activity (**Fig 4E**).

## Discussion

Two concentric cylinders of stem cells function in the root to accommodate the vast increase in girth during secondary growth: the vascular and cork cambium (**Fig S1A**). Each of these stem cell regions can produce cells towards the inside or the outside, and the relative amounts depend on the phase of development but can be influenced by environmental factors^9,10^. Our results shed light on the molecular regulation of cambium activity by showing that the phosphorylation status of ANT controls this differential output of both cambia (**Fig 4E**). ANT is phosphorylated via the ER-MPK6 receptor pathway, and this is promoted by local auxin signalling. As such, this auxin-dependent ANT phospho-switch defines the directionality of the vascular and cork cambium activity and thus controls the relative proportions of cell types being made during secondary growth.

What is known to date is that, during plant growth, the ERECTA pathway acts by phosphorylating WRKY TFs during embryogenesis^35,51,52^, and *bHLH* TFs during stomata formation^39,43,53–56^, and *AP2* factors during radial growth. This suggests that while the ER-MPK6 receptor pathway is conserved, developmental specificity is achieved at the level of the TF family that is phosphorylated. However, *AP2* family TFs such as *BABY BOOM* / *PLETHORA4* (*AP2* family) and *bHLH* family TFs: *TMO5*, *LHW*, and *T5Ls* (*bHLH* family), which are known to regulate cell proliferation during embryogenesis^57^ and vascular development^58,59^, respectively, is regulated by ER-mediated phosphorylation. An alternative explanation could, therefore, be that only a few ER targets have been identified and that ER-modulated processes share a similar downstream TF module. Future research will show if ER-controlled processes are governed by common core modules of transcriptional regulators of the *AP2*, *bHLH*, and *WRKY* families.

Interestingly, a recent study revealed complex regulation of vascular development through the LRR-RLK receptor PXY by its ligand TDIF. Activation of this pathway promotes vascular cambium proliferation via *ANT*/*AIL* (AINTEGUMENTA-like) transcription factors such as *AIL5*/*PLT5* (*PLETHORA 5*), which inhibits xylem vessel and phloem sieve element formation^33^. Furthermore, it has been recently shown that ER forms a complex with PXY, which is needed for PXY-TDIF responses during xylem differentiation^60^, raising the question of whether ANT phosphorylation requires ER-PXY complex formation and whether other members of the ANT family are phosphorylated. Interestingly, among the genes downregulated upon blocking auxin signalling in the cambia, we found *AIL5* and *AIL7* (**Fig S3C**). Therefore, future research focusing on the interaction of ER with all members of the PXY family (*PXY* is not expressed in the cork cambium^7^) and whether activation of ER/PXY signalling triggers ANT/AILs phosphorylation are necessary to dissect the different contributions and specificity of LRR-RLK receptors in balancing cambia output by ANT/AILs phospho-switch.

Altogether, these results place ANT at the cross-talk of several pathways, highlighting the possibility that *ANT* transcript levels and phosphorylation status can be exploited to boost root radial growth and favour the formation of cell types that are relevant for biomass accumulation, such as xylem and cork (**Fig S10A-C**). This will be valuable in tree breeding programs aiming to enforce biomass accumulation as wood^61^ or in initiatives aiming to store more carbons in the cork and xylem of roots^1,2,62^, which represent strategies to mitigate the effects of global warming.

## Materials and Methods

### Plant material and growth

All Arabidopsis mutants and transgenic lines (T1 segregating, F1 crossing, T2 segregating and/or T3 homozygous populations) used in this study are described in **Supplementary Table 1**. The plants were grown on 1/2MS plates with 1% sugar and 0.8% plant agar. DEX (Dexamethasone) treatment was performed by transferring plants on plates containing 10µM DEX (Sigma, D1756). Auxin treatment was performed by transferring plants to plates containing 1µM NAA (1-naphthaleneacetic acid) plates (1µM, Duchefa, N0903). The Arabidopsis plants were grown *in vitro* under continuous light until sampling, or in soil under long-day conditions (16 hours light versus 8 hours dark) at 22 Celsius degrees.

### Molecular cloning

Promoters were amplified from *Arabidopsis thaliana* genomic DNA and protein coding sequence from root cDNA with the primers listed i**n Supplementary Table 2** and cloned into Green Gate entry vectors^63^. All final constructs were assembled with the modular green gate technology^63^. If promoters or protein coding sequences contain BSA I sites, they were replaced as described in^64^. All the modules and vectors used in this study are listed in **Supplementary Table 3**. In **Supplementary Table 4**, all the module assembly strategies are described. All final vectors were transformed into *Agrobacterium tumefaciens* GV3101, and *Arabidopsis thaliana* plants were transformed via floral dipping. F1 (from crossing) seeds, T1 and T2 seeds selected via fast-red fluorescence seed selection^65^ or Homozygous T3 lines were used in this study.

### Histology and fluorescent staining

0.5 cm root segments have been collected for plastic embedding (at 0.5 cm below the root/hypocotyl junction). Plastic embedding was performed in Technovit 7100 (Heraeus Kulzer; 64709003) following the manufacturer’s instructions and as described in^66^. The plastic sections (5-8 cm) were stained with 0.1 % toluidine blue (Sigma; T3260). The pictures were acquired with a Zeiss Axio M2 imager microscope or a Zeiss Axiophot microscope, with the ZEN Blue software (version 3.4). Suberin was stained by Fluorol yellow (FY), followed the instruction in^67^, using Fluorol yellow (Santa Cruz, sc-215052). Propidium Iodide (PI) staining was achieved by directly mounting the root in a 10-30µg/ml solution (Sigma, P4864).

### Confocal Microscopy

A Zeiss confocal (LSM880/LSM710) was used to acquire the confocal images with the following settings. For GFP (green fluorescent protein): ex. 488 nm; em. 485–505 nm. For YFP (yellow fluorescent protein), mCitrine and Venus: ex. 514 nm; em. 515–545 nm. For mCherry, RFP, and PI: ex. 561 nm; em. 570–630 nm. For FY: ex. 488nm; em 480-550nm. Orthogonal views of a Z stack were obtained using the ZEN Black software (Zen 2.3 SP1).

### RNA sequencing

RNA was extracted from the uppermost 1.5 cm of roots (12-day-old plants) for each sample (Mock or DEX treatment; approximately 100 plants for each sample), using a Universal RNA Purification Kit (Roboklon, E3598-02). RNA quality was checked by Agilent RNA Bioanalyzer chip traces and RNA quantity was estimated using a Nanodrop 2000.

The enrichment of mRNA by oligo-dT pull-down was performed using the NEBNext Poly(A) mRNA Magnetic Isolation Module (NEB #E7490L). The construction of RNA libraries was carried out using the NEBNext Ultra II RNA Library Prep Kit for Illumina (NEB #E7530L) with NEBNext Multiplex Oligos for Illumina (Index Primers Set 1 to Set 4). Finally, the RNA library size and quality were measured on DNA High Sensitivity Bioanalyzer chip (Agilent), and RNA libraries were quantified by the NEBNext Library Quant Kit for Illumina (NEB #E7630S).

The pair-end 150bp RNA sequencing was performed by Novogene. The analysis of the data was performed using the Galaxy platform (https://usegalaxy.eu/)^68^. Adaptors were removed with Trimmomatic using default parameters, and read quality was assessed with FastQC. Reads were aligned to the TAIR10 genome using Salmon quant using default parameters. Reads were counted using Tximport, and differential gene expression analysis was performed using DESeq2 (**DataS1)**.

### Statistical analyses

Quantification, statistical analysis and plotting were performed using in R (version 4.0.3). All box plots were generated using ggplot2 (version 3.4.3). Count data was analysed by fitting generalized linear (mixed) models following poisson (stats package, version 4.0.3), quasi-poisson (stats package, version 4.0.3), or negative binomial (stats package, version 4.0.3). Residual diagnostics of the models were performed using the simulation-based approach of the DHARMa package (version 0.4.6) to select the best fitting model. Inference from the models was done with the emmeans package (version 1.5.5-1) and p-values for the pairwise contrasts were adjusted using the Holm-Bonferroni method. Compact letter display of pairwise comparisons was generated using the multcomp package (version 1.4-16), using a significance level of 0.05. All data and tests used for statistical analyses are summarised in Table S2.

### Cell number quantification, root area, and image analyses

The identification of periderm and secondary vascular tissues was based on a group of specialized cells named phloem pools (red asterisks), which demarcated the boundary between these tissue systems, serving as persistent histological markers before, at, and after the secondary growth **(Fig S1A-B)_6,7_.** The quantification of periderm layers was described in^7^. The cell number quantification was performed by Fiji (https://fiji.sc/)^69^. For every genotype, 8-25 cross-sections from independent plants were analysed. 2-3 independent experiments were performed and only one experiment was presented. The cork/xylem/periderm/vasculature tissue cell number was calculated using the cell counter plugin in Fiji. To identify specific cell types within secondary vascular tissues, the p*PXY*::er-CFP marker was introduced into several transgenic lines and this fluorescent marker served as a reliable indicator for distinguishing different cell populations. In vibratome cross-sections, the quantification of xylem cell number is based on secondary cell wall accumulation, the quantification of vascular cambium cells by expression of the p*PXY*::er-CFP marker, and the quantification of phloem cells is calculated by subtraction (the number of secondary growth tissues cells minus the number of xylem and vascular cambium cells). The tissue cell numbers were calculated using the cell counter plugin in Fiji. The area (in pixels) of the root was calculated in Fiji.

### Western blot and phosphorylation assay in *Nicotiana benthamiana* leaves

Using the GreenGate cloning system, all CDS were driven by 35S promoter and tagged with HA-tag or FLAG-tag at the C-terminus (details in Table S4). All plant binary vectors were transformed into *Agrobacterium tumefaciens* (strain GV3101). These vectors were cultured overnight and resuspended in the tobacco infiltration buffer [10 mM MES (pH 5.7) and 10 mM MgCl_2_] for each construct. The selected constructs were co-infiltered with the P19 silencing vector into the leaves of *N. benthamiana* and the infiltrated leaves were collected after 72h. The leaves samples were ground to fine powders in liquid nitrogen, and total proteins were extracted in SDS-loading buffer [62.5 mM Tris-HCl pH 6.8, 2.5 % SDS, 0.002 % Bromophenol Blue, 0.7135 M (5%) DTT, 10 % glycerol, and 12.5 mM Protease Inhibitor Cocktail] followed by heating at 95 °C for 5 min. Calf-intestinal alkaline phosphatase (CIP) (NEB #M0525V) was added to protein samples and incubated at 37°C for 1h. Protein samples were resolved in a Phos-tag (AAL-107, Wako Pure Chemical Corporation) 10% SDS-PAGE gel and analysed by Western blot.

The primary antibody Anti-c-HA (Abcam, ab18181, 1:2,500 dilution) and Anti-FLAG Nanobody-HRP (AlpalifebioK, TSM1318, 1:5,000 dilution) and secondary antibody GOAT ANTI MOUSE (Bio-rad, STAR137, 1:20,000 dilution) were selected for the immunoblotting. Signal detection was performed with the Clarity Western ECL Substrate (Bio-rad, 170-5060) and detected by ChemiDoc XRS+ System.

### Vibratome sections for confocal imaging

The uppermost 1 cm of roots were collected and fixed in 4% paraformaldehyde (PFA) in Ca^2+^, Mg^2+^-free PBS. The samples were vacuum infiltrated at RT for 1h. After washing with PBS, samples were embedded in warm 5% agarose (Sigma-Aldrich, A4018). Once the agarose solidified, the blocks were sectioned using a vibratome (Leica, VT1000 S) at a thickness of 80-100 µm per section. The sections were washed 3 times with PBS and stained with 1μg/ml Calcofluor White (Merck, 18909) or 0.05% Direct Red 23 (Sigma, 212490).

### Edu staining

The plants were grown on 1/2 MS plates for 17d. Next, plants were collected together with some agar, and transferred into liquid 10µM EdU (Sigma, BCK-EDU488) 1/2 MS medium for 16h. The samples were then fixed and sectioned as described in vibratome confocal section above. The cross-sections were collected and submerged by Edu detection cocktail under RT for 1h following manufacturer instructions. The cross-sections were washed 2 times with PBS and stained with 1μg/ml Calcofluor White.

## Acknowledgments

We thank Ari Pekka Mähönen and Peter Etchells for sharing seeds and materials. We thank Andrea Boch and Xudong Zhang for technical help. We thank Perrine Dalby for the initial experiment on the mutants of *ER*. We thank Shihao Su for critical reading of the manuscript. We thank Max Minne and Nataliia Konstantinova for their help with statistical analysis.

## Author contributions

W.X, and L.R, conceived the project. W.X, L.Y, M.B and L.R, designed the experiments. W.X, and L.Y, conducted the experiments, and analysed the data. W.X. performed RNA-seq, and W.X, and L.R, analysed the obtained data. W.X, L.Y, H.C, D.M, and Y.M, generated the plant transgenic lines. W.X, H.J. and L.R, acquired the confocal images. W.X, performed the qRT-PCRs and plant embedding. W.X, L.Y, and H.J, cross the plants. W.X, L.Y, and S.Y, performed the phosphorylation experiments. W.X, and L.Y, performed the western blot. W.X, analysed data measurement. W.X., B.D.R. and LR wrote the paper with input from all authors.

## Data and materials availability

Raw RNA-seq data can be accessed at NCBI with GEO number GSE270329.

## Supplementary Materials

Figs S1 to S10

Tables S1 to S5

Data S1 to S2

Supplementary references (1-14)

## Supplementary Figure Legends

**Fig. S1. Vascular cambium and cork cambium dynamics during secondary growth. Related to Figure 1**.

**(A)**, Sketches highlighting the different tissues occurring during secondary growth. In the Middle panel, secondary vascular (SV) tissues are highlighted in orange, and the periderm in pink. Left panels: tissues comprising the secondary vasculature tissues: xylem (pale blue), vascular cambium (brown) and phloem (violet). Right panels: tissues comprising the periderm: cork (military green), cork cambium (green) and phelloderm (turquoise). **(B)**, Cross-sections highlighting how the different tissues were morphologically identified and scored. The blue line shows the boundary between the secondary vascular tissues and the periderm. Yellow asterisks indicate pericycle cells and red asterisks indicate the phloem poles. **(C)**, Left panels: vibratome-sections of the uppermost part of p*PXY*::*shy2-2*-GR x p*PXY*::er-CFP F1 plants. 12-day-old plants were transferred to either Mock (left-up panel) or 10 μM DEX (left-down panel) plates for 2 days. Right panel: quantification of the number of xylem cells, vascular cambium cells and phloem cells in left panels. White asterisk indicates xylem cells. **(D)**, Left panels: cross-sections of the most mature part of 12-day-old and 20-day-old roots with highlighted the periderm development. The orange squares indicate the magnification area shown in the middle panels (middle panels). Black dots represent periderm cell layers. Right panels: cross-sections (plastic embedding) stained with Fluorol yellow (FY) of the uppermost part of 12-day-old and 20-day-old roots. Yellow dots represent suberized periderm layers and grey dots represent non-suberized periderm layers. **(E)**, Quantification of periderm layers number of the experiment shown in (FigS1D). **(F)**, Quantification of the number of periderm and cork cells in (FigS1D). Black and white scale bars: 20μm. Count data was modelled with a generalized mixed model. Here the Poisson mix model was selected. P values were adjusted using Holm-Bonferroni correction.

**Fig. S2. Auxin balances the formation of the cork cambium derivatives. Related to Figure 1**.

**(A)**, Cross-sections (plastic embedding) of the uppermost part of Col0 roots. 12-day-old plants were transferred for 2 days on Mock (left panel) or 1μM NAA (right panel) plates. Black dots represent periderm cell layers. **(B)**, Cross-sections of the upper most part of Col0 roots stained with Fluorol yellow (FY). 12-day-old plants were transferred for 2 days on Mock (left panel) or 1μM NAA (right panel) plates. Yellow dots represent suberized periderm layers and grey dots represent non-suberized periderm layers. **(C)**, Quantification of periderm layers of the experiment shown in (Fig S2A). **(D)**, Quantification of the number of periderm and cork in (FigS2A). **(E)**, Cross-sections (plastic embedding) quantification of the uppermost part of 20-day-old p*PER15*::*slr*-GR (p*MYB84*::NLS-3xGFP and W131Y background) roots; 12-day-old plants were transferred for 8 days on Mock (left panel) or 10μM DEX (right panel) plates. **(F)**, Cross-sections of the uppermost part of p*PER15*::*slr*-GR roots with Fluorol yellow (FY). 12-day-old plants were transferred for 8 days on Mock (left panel) or 10μM DEX (right panel) plates. Yellow dots represent suberized periderm layers and grey dots represent non-suberized periderm layers. **(G)**, Quantification of periderm layers of the experiment shown in (FigS2E). **(H)**, Quantification of the number of periderm and cork cells in (FigS2E). Black and white scale bars: 20μm. Count data was modelled with a generalized mixed model. Here the Poisson mix model was selected. P values were adjusted using Holm-Bonferroni correction.

**Fig. S3. ER is induced by auxin and promotes secondary growth. Related to Figure 2 and data S1.**

**(A)**, The sketch of the RNA-seq experimental design. 12-day-old plants were transferred for 24h on Mock or 10μM DEX plates, and the uppermost 1.5 cm-2cm was collected for each sample. **(B)**, Venn diagrams show the overlap of genes co-upregulated or genes co-downregulated in p*PER15*::*slr*-GR and p*PXY*::*shy2-2*-GR in Mock or 10μM DEX treatment (log2FC≥0.5 or log2FC≤-0.5; and padj≤0.05). **(C)**, The heatmap of genes regulated by DEX treatment in both p*PER15*::*slr*-GR and p*PXY*::*shy2-2*-GR. **(D)**, Cross-sections (plastic embedding) of the uppermost part of Col0, *erl1*, *erl2* and *er* roots. 6-day-old plants were transferred for 2 days on Mock (left panels) or 1μM NAA (right panels) plates. **(E)**, Quantification of the number of secondary vascular (SV) tissues and xylem cells in (FigS3D). M indicates Mock and N indicates NAA. **(F)**, Cross-sections (plastic embedding) of the uppermost part of 12-day-old p*ER*::ER-YPet, *er erl1 erl2* (left panel), and p*PXY*::NSImb-vhhGFP4 in p*ER*::ER-YPet*, er erl1 erl2* (right panel) roots. The xylem vessels are highlighted in blue. **(G)**, Quantification of the number of secondary vascular tissues and xylem cells in (FigS3F). Black scale bars: 20μm. Count data was modelled with a generalized mixed model. Here the Poisson mix model was selected. P values were adjusted using Holm-Bonferroni correction.

**Fig. S4. ER controls vascular cambium derivative formation. Related to Figure 2**.

**(A)**, Relative expression of *ER*, *ERL1* and *ERL2* in the Col0. 12-day-old plants were transferred for 1 days on Mock or 1μM NAA plates. **(B)**, Orthogonal view of z stacks of p*ER*::3xVenus-N7. 12-day-old roots at the positions corresponding to stage 1 (lower panel, before the initiation of periderm), stages 3 (middle panel, on the initiation of periderm), and stage 5 (up panel, after the initiation of periderm) of secondary growth. The red arrow indicates cork cambium/pericycle and the green arrow indicates vascular cambium. PI: Propidium Iodide. **(C)**, Cross-sections (plastic embedding) of the uppermost part of 20-day-old p*PXY*::BIR3-ER (right panel) roots. The blue arrow indicates the ectopic secondary xylem cells. The xylem vessels are highlighted in blue. **(D)**, Upper panels: cross-sections (plastic embedding) of the uppermost part of 13-day-old Col0 and p*SMXL5*::BIR3-ER. Lower panel: quantification of the number of secondary vascular (SV) tissues and xylem cells in (FigS4D). **(E),** Vibratome-sections of the uppermost part of 20-day-old Col0 x p*PXY*::er-CFP (up-panel), and p*PXY*::BIR3-ER x p*PXY*::er-CFP (low-panel) F1 roots. White Asterisk indicates xylem cells. **(F),** Quantification of the number of xylem cells, vascular cambium cells and Phloem cells in (FigS4E). Black and white scale bars: 20μm. Count data was modelled with a generalized mixed model. Here the Poisson mix model was selected. P values were adjusted using Holm-Bonferroni correction.

**Fig. S5. ANT is phosphorylated by MPK6. Related to Figure 2 and Figure 3.**

**(A)**, Relative expression of *BP*, *WOX4*, and *ANT* in the Col0 and p*PXY*::BIR3-ER roots. 14-day-old roots for collection. **(B)**, Orthogonal view of Z-stacks of p*ANT*::H2B-YFP. 12-day-old roots at the positions corresponding to stage 1 (lower panel), stages 3 (middle panel), and stage 5 (up panel) of secondary growth. The red arrow indicates the cork cambium/pericycle and the green arrow indicates the vascular cambium**. (C),** Upper panels: orthogonal view of Z-stacks of p*MPK6*::NLS-3xmVenus. 14-day-old roots at the positions corresponding to stage 1 (lower panel), stages 3 (middle panel), and stage 5 (up panel) of secondary growth. Lower panels: vibratome-sections of the uppermost part of 15-day-old p*MPK6*::NLS-3xmVenus. The red arrow indicates the cork cambium / pericycle and the green arrow indicates the vascular cambium. PI: Propidium Iodide; DR: Direct Red. **(D)**, Putative MPK6 phosphorylation sites in the ANT sequence. ANT contains two AP2 domains. Four putative MPK6 phosphorylation residues (S113, S202, T496 and S526) were highlighted in the ANT protein. **(E)**, *In vivo* phosphorylation assays in *N. benthamiana.* The p*35S*::ANT-4D-3xFLAG (Phosphor-mimic version of ANT), the p*35S*::ANT-4A-3xFLAG (Phospho-dead version of ANT) and the p*35S*::ANT-3xFLAG vectors were transiently expressed in leaves. Protein extracts were analysed on a Phos-tag/Normal SDS-PAGE gel and probed with an anti-Flag antibody. Black scale bars: 20μm. Count data was modelled with a generalized mixed model. Here the Poisson mix model or Poisson model was selected. P values were adjusted using Holm-Bonferroni correction.

**Fig. S6. ANT acts downstream ER signalling to control vascular cambium output. Related to Figure 3**.

**(A),** Vibratome-sections of the uppermost part of p*PXY*::ANT-GR x p*PXY*::er-CFP F1 plants. 7-day-old plants were transferred to either Mock (left panel) or 10 μM DEX (right panel) plates for 5 days. White asterisk indicates xylem cells. **(B),** Quantification of the number of xylem cells, vascular cambium cells and phloem cells in (FigS6A). **(C),** Cross-sections (plastic embedding) of the uppermost part of 14-day-old Col0, and T1 of p*PXY*::ANT-GFP and p*PXY*::ANT-4A-GFP roots. The xylem vessels are highlighted in blue. **(D),** Quantification of xylem cell number and secondary vascular (SV) tissues cell number in 14-day-old root shown in (FigS6C). **(E),** Vibratome-sections of the uppermost part of p*PXY*::ANT-SRDX-GR x p*PXY*::er-CFP F1 plants. 7-day-old plants were transferred to either Mock (left panel) or 10 μM DEX (right panel) plates for 5 days. White asterisk indicates xylem cells. **(F),** Quantification of the number of xylem cells, vascular cambium cells and phloem cells in (FigS6E). Black and white scale bars: 20μm. Count data was modelled with a generalized mixed model. Here the Poisson mix model or Poisson model was selected. P values were adjusted using Holm-Bonferroni correction. Count data was also analysed using the Student’s T-test.

**Fig. S7. ANT phosphorylation depends on ER signalling. Related to Figure 3**.

**(A),** Upper panels: root cross-sections of p*PXY*::BIR3-ER x p*PXY*::ANT-SRDX-GR. 12-day-old plants were transferred to either Mock (left panel) or 10 μM DEX (right panel) plates for 8 days. Lower panel: quantification of 20-day-old root in xylem cell number and secondary vascular tissues cell number of the experiment shown in (FigS7A). **(B-D),** The root cross-sections of T1 p*PXY*::ANT-GR (B), p*PXY*::ANT-4A-GR (C), and p*PXY*::ANT-4D-GR (D) in p*PXY*::NSImb-vhhGFP4 in p*ER*::ER-YPet*, er erl1 erl2* plants. 12-day-old plants were grown on Mock (left panel) or 10μM DEX (right panel) plates. The xylem vessels are highlighted in blue. **(E),** Quantification of 12-day-old root in xylem cell number and secondary vascular tissues cell number of the experiment shown in (FigS7B-D). Black scale bars: 20μm. Count data was modelled with a generalized mixed model. Here the Poisson mix model or Poisson model was selected. P values were adjusted using Holm-Bonferroni correction.

**Fig. S8. ER balances cork cambium produces. Related to Figure 4**.

**(A)**, Cross-sections of the uppermost part of 12-day-old Col0 (left panel) and p*PER15*::BIR3-ER (right panel) roots. Black dots represent periderm cell layers. **(B)**, Cross-sections (plastic embedding) stained with Fluorol yellow (FY) of the uppermost part of 12-day-old Col0 (left panel) and p*PER15*::BIR3-ER (right panel) roots. Yellow dots represent suberized periderm layers and grey dots represent non-suberized periderm layers. **(C)**, Quantification of periderm layers number of the experiment shown in (Fig S8A). **(D)**, Quantification of the number of periderm and cork cells in (FigS8A). **(E)**, Cross-sections of the uppermost part of 12-day-old p*ER*::ER-YPet, *er erl1 erl2* (left panel), p*PER15*::NSImb-vhhGFP4 in p*ER*::ER-YPet*, er erl1 erl2* (right panel) roots. Black dots represent periderm cell layers. **(F)**, Cross-sections (plastic embedding) stained with Fluorol yellow (FY) of the uppermost part of 12-day-old Col0 (left panel) and p*PER15*::BIR3-ER (right panel) roots. Yellow dots represent suberized periderm layers and grey dots represent non-suberized periderm layers. **(G)**, Quantification of periderm layers of the experiment shown in (Fig S8E). **(H)**, Quantification of the number of periderm and cork cells in (FigS8E). White scale bars: 20μm. Count data was modelled with a generalized mixed model. Here the Poisson mix model or Poisson model was selected. P values were adjusted using Holm-Bonferroni correction.

**Fig. S9. ANT balances cork cambium derivative formation. Related to Figure 4**.

**(A)**, Cross-sections of the uppermost part of 12-day-old p*PER15*::ANT-GR roots; 7-day-old plants were transferred for 5 days on Mock (left panel) or 10μM DEX (right panel) plates. Black dots represent periderm cell layers. **(B)**, Cross-sections (plastic embedding) stained with Fluorol yellow (FY) of the uppermost part of 12-day-old p*PER15*::ANT-GR roots. 7-day-old plants were transferred for 5 days on Mock (left panel) or 10μM DEX (right panel) plates. Yellow dots represent suberized periderm layers and grey dots represent non-suberized periderm layers. **(C)**, Quantification of periderm layers of the experiment shown in (FigS9A). **(D)**, Quantification of the number of periderm and cork in (FigS9A). White scale bars: 20μm. Count data was modelled with a generalized mixed model. Here the Poisson mix model was selected. P values were adjusted using Holm-Bonferroni correction.

**Fig. S10. Extra secondary xylem cells enhance the plant growth. Related to Figure 4. (A)**, Root cross-section of p*PXY*::ANT-SRDX-GR plants grown in Mock/DEX plates for 20 day are shown. 12-day-old plants on Mock (up panel) or DEX (lower panel) plates and were transferred to Mock plates. **(B-C)**, Quantification of fresh weight of the shoot **(B)** and area of the root **(C)** is measured in the experiment shown in (FigS10A). White scale bars: 20μm. Count data was modelled with a generalized mixed model. Here the Poisson mix model was selected. P values were adjusted using Holm-Bonferroni correction.

