## supplemental Files for "An AINTEGUMENTA phospho-switch controls bilateral stem cell activity during secondary growth"

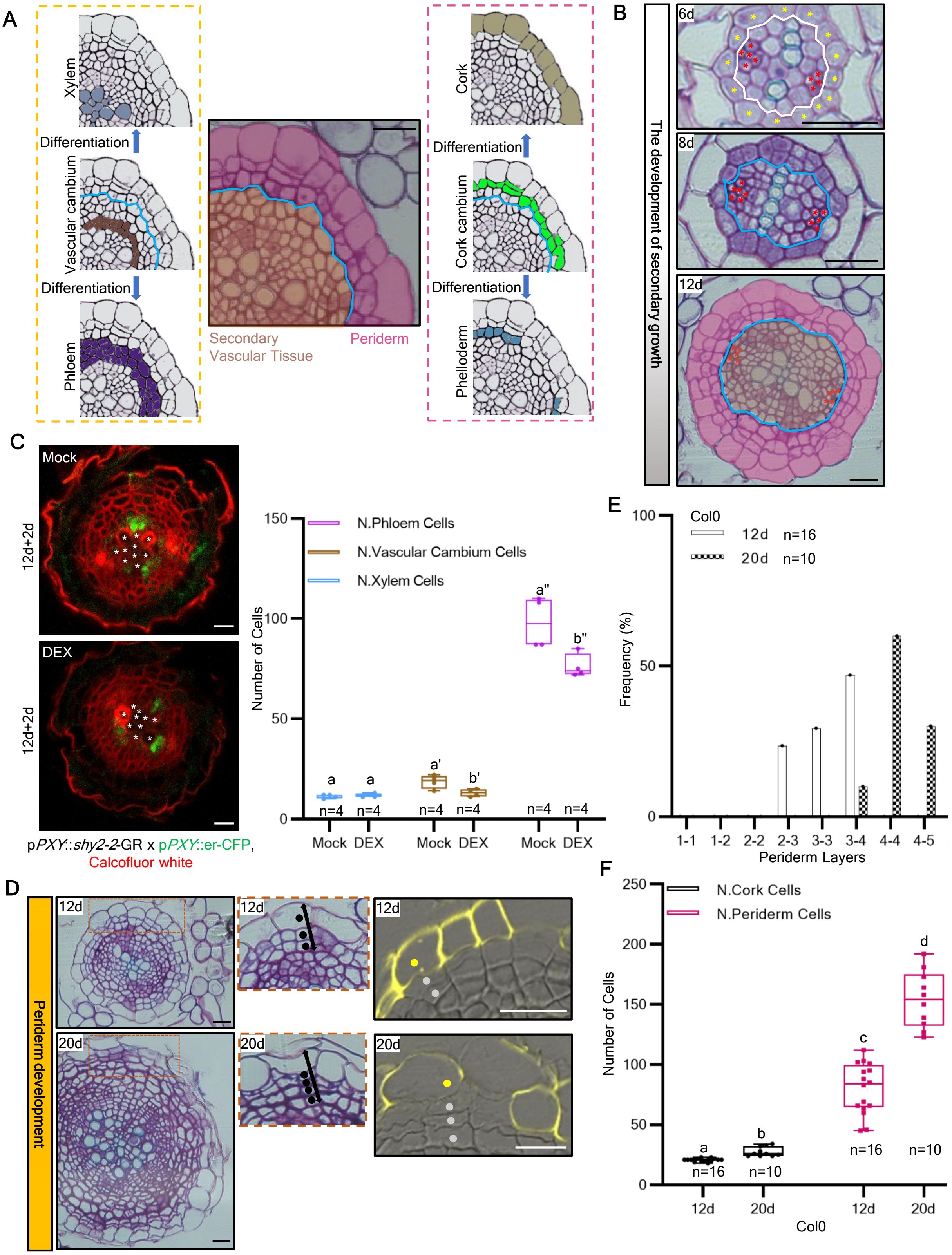
A
Xylem
Differentiation
Vascular cambium
Differentiation
Phloem
Secondary
Vascular Tissue
Periderm
Cork
Differentiation
Cork cambium
Differentiation
Phelloderm
B
The development of secondary growth
6d
8d
12d
C
Mock
DEX
12d+2d
pPXY::shy2-2-GR x pPXY::er-CFP,
Calcofluor white
N.Phloem Cells
N.Vascular Cambium Cells
N.Xylem Cells
Number of Cells
Mock DEX Mock DEX Mock DEX
n=4
D
Periderm development
12d
20d
E
Col0
12d n=16
20d n=10
Frequency (%)
Periderm Layers
1-1 1-2 2-2 2-3 3-3 3-4 4-4 4-5
F
N.Cork Cells
N.Periderm Cells
Number of Cells
n=16 n=10
12d 20d 12d 20d
Col0

**Fig. S1. Vascular cambium and cork cambium dynamics during secondary growth. Related to Figure 1.**

**(A)**, Sketches highlighting the different tissues occurring during secondary growth. In the Middle panel, secondary vascular (SV) tissues are highlighted in orange, and the periderm in pink. Left panels: tissues comprising the secondary vasculature tissues: xylem (pale blue), vascular cambium (brown) and phloem (violet). Right panels: tissues comprising the periderm: cork (military green), cork cambium (green) and phelloderm (turquoise). **(B)**, Cross-sections highlighting how the different tissues were morphologically identified and scored. The blue line shows the boundary between the secondary vascular tissues and the periderm. Yellow asterisks indicate pericycle cells and red asterisks indicate the phloem poles. **(C)**, Left panels: vibratome-sections of the uppermost part of p*PXY*::*shy2-2*-GR x p*PXY*::ER-CFP F1 plants. 12-day-old plants were transferred to either Mock (left-up panel) or 10 µM DEX (left-down panel) plates for 2 days. Right panel: quantification of the number of xylem cells, vascular cambium cells and phloem cells in left panels. White asterisk indicates xylem cells. **(D)**, Left panels: cross-sections of the most mature part of 12-day-old and 20-day-old roots with highlighted the periderm development. The orange squares indicate the magnification area shown in the middle panels (middle panels). Black dots represent periderm cell layers. Right panels: cross-sections (plastic embedding) stained with Fluorol yellow (FY) of the uppermost part of 12-day-old and 20-day-old roots. Yellow dots represent suberized periderm layers and grey dots represent non-suberized periderm layers. **(E)**, Quantification of periderm layers number of the experiment shown in (FigS1D). **(F)**, Quantification of the number of periderm and cork cells in (FigS1D). Black and white scale bars: 20µm. Count data was modelled with a generalized mixed model. Here the Poisson mix model was selected. P values were adjusted using Holm-Bonferroni correction.

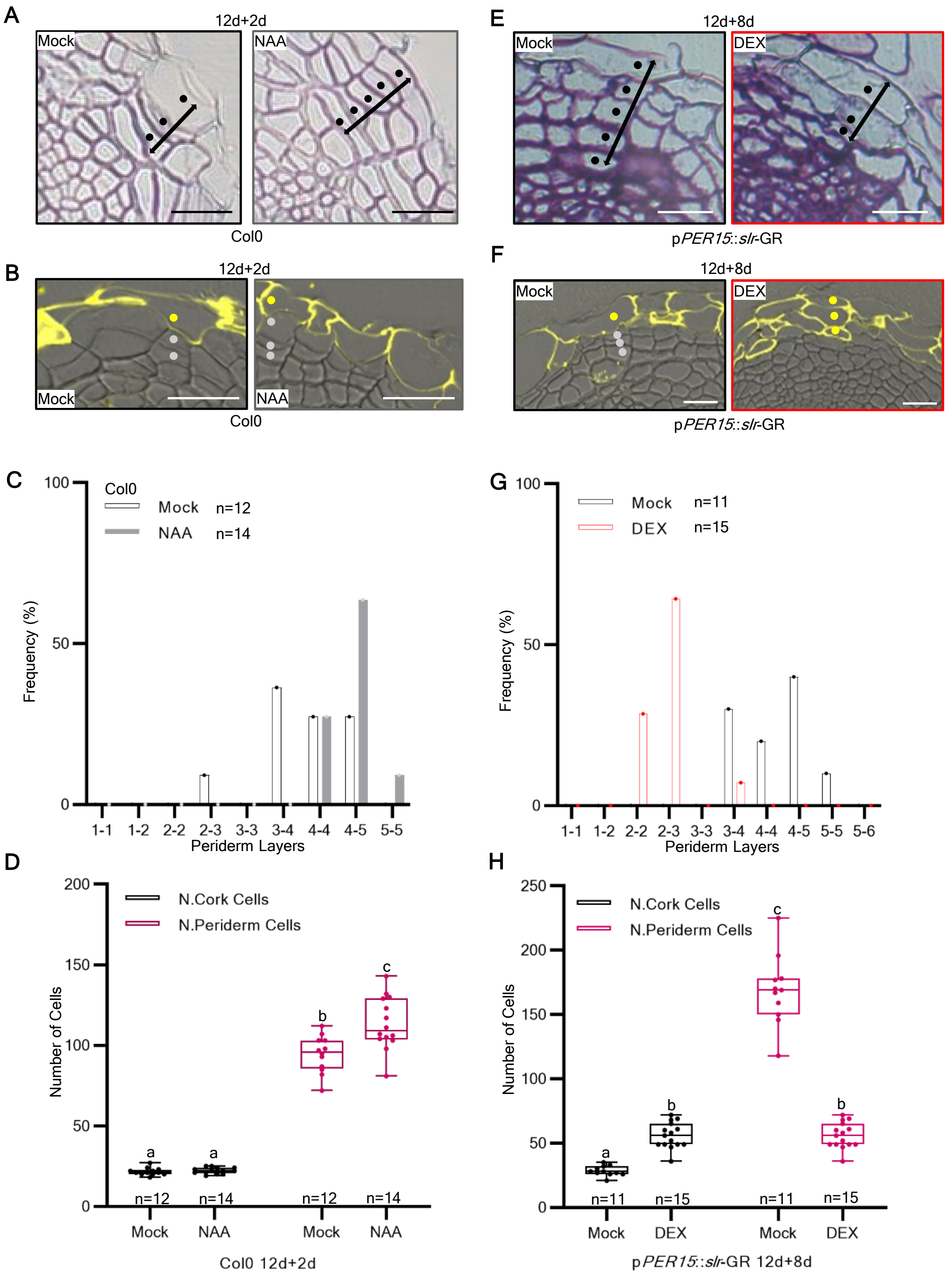


**Fig. S2. Auxin balances the formation of the different cork cambium derivatives. Related to Figure 1.**
**(A)**, Cross-sections (plastic embedding) of the uppermost part of Col0 roots. 12-day-old plants were transferred for 2 days on Mock (left panel) or 1µM NAA (right panel) plates. Black dots represent periderm cell layers. **(B)**, Cross-sections of the upper most part of Col0 roots stained with Fluorol yellow (FY). 12-day-old plants were transferred for 2 days on Mock (left panel) or 1µM NAA (right panel) plates. Yellow dots represent suberized periderm layers and grey dots represent non-suberized periderm layers. **(C)**, Quantification of periderm layers of the experiment shown in (Fig S2A). **(D)**, Quantification of the number of periderm and cork in (FigS2A). **(E)**, Cross-sections (plastic embedding) quantification of the uppermost part of 20-day-old p*PER15*::*slr*-GR (p*MYB84*::NLS-3xGFP and W131Y background) roots; 12-day-old plants were transferred for 8 days on Mock (left panel) or 10µM DEX (right panel) plates. **(F)**, Cross-sections of the uppermost part of p*PER15*::*slr*-GR roots with Fluorol yellow (FY). 12-day-old plants were transferred for 8 days on Mock (left panel) or 10µM DEX (right panel) plates. Yellow dots represent suberized periderm layers and grey dots represent non-suberized periderm layers. **(G)**, Quantification of periderm layers of the experiment shown in (FigS2E). **(H)**, Quantification of the number of periderm and cork cells in (FigS2E). Black and white scale bars: 20µm. Count data was modelled with a generalized mixed model. Here the Poisson mix model was selected. P values were adjusted using Holm-Bonferroni correction.

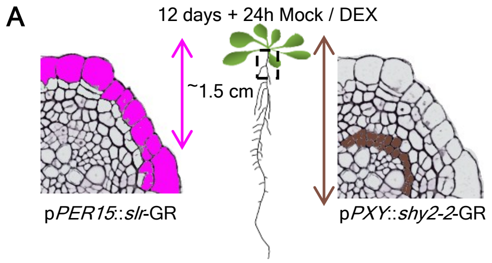


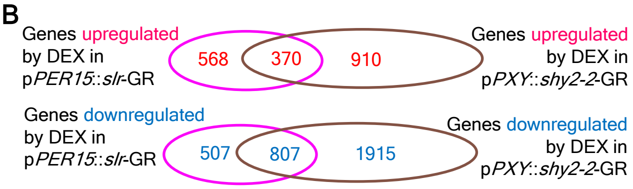


C

| Gene | $Log_2FC$ (p*PER15*::*slr*-GR / p*PXY*::*shy2-2*-GR) | P-value (p*PER15*::*slr*-GR / p*PXY*::*shy2-2*-GR) |
|---|---|---|
| *BP* (*AT4G08150*) | -0.76 / -0.93 | P<0.05 / P<0.05 |
| *WOX4* (*AT1G46480*) | -1.43 / -1.70 | P<0.05 / P<0.05 |
| *ANT* (*AT4G37750*) | -1.24 / -3.48 | P<0.05 / P<0.05 |
| *ER* (*AT2G26330*) | -1.36 / -1.41 | P<0.05 / P<0.05 |
| *LBD3* (*AT1G16530*) | -0.66 / -0.86 | P<0.05 / P<0.05 |
| *PIN4* (*AT2G01420*) | -1.01 / -0.69 | P<0.05 / P<0.05 |
| *WOX14* (*AT1G20700*) | -0.63 / -0.75 | P<0.05 / P<0.05 |
| *ERL1* (*AT5G62230*) | -1.32 / 6.73 | ns / ns |
| *ERL2* (*AT5G62230*) | 0.00 / 6.38 | ns / ns |
| *PLT1* (*AT3G20840*) | -0.28 / -0.02 | ns / ns |
| *PLT2* (*AT1G51190*) | -0.29 / 0.23 | ns / ns |
| *AIL6/PLT3* (*AT5G10510*) | -0.27 / 0.04 | ns / ns |
| *AIL5/PLT5* (*AT5G57390*) | -0.29 / -1.70 | ns / 0.00 |
| *AIL7/PLT7* (*AT5G65510*) | -1.16 / -0.17 | 0.00 / ns |

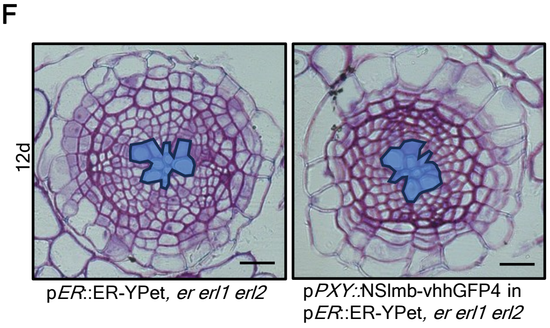


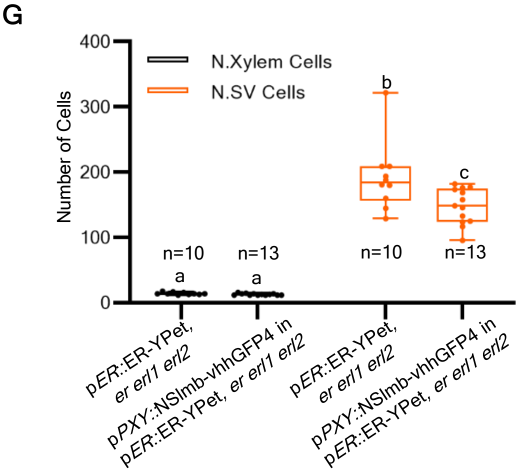


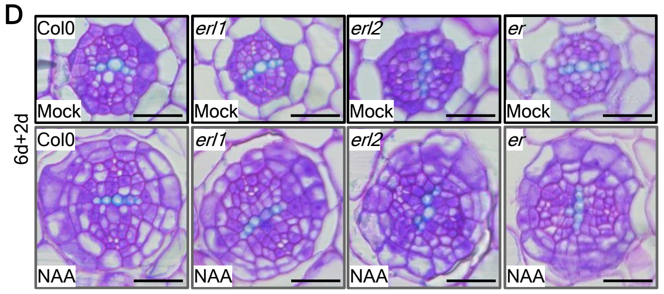


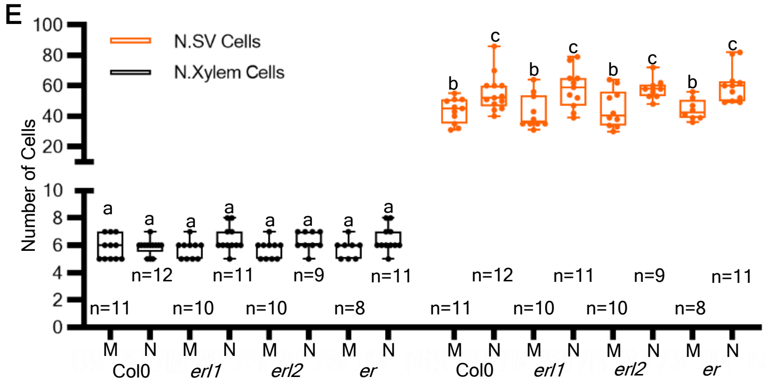


**Fig. S3. ER is induced by auxin and promotes secondary growth. Related to Figure 2 and data S1.**
**(A)**, The sketch of the RNA-seq experimental design. 12-day-old plants were transferred for 24h on Mock or 10µM DEX plates, and the uppermost 1.5 cm- 2cm was collected for each sample. **(B)**, Venn diagrams show the overlap of genes co-upregulated or genes co-downregulated in p*PER15*::*slr*-GR and p*PXY*::*shy2-2*-GR in Mock or 10µM DEX treatment (log2FC≥0.5 or log2FC≤-0.5; and padj≤0.05). **(C)**, The heatmap of genes regulated by DEX treatment in both p*PER15*::*slr*-GR and p*PXY*::*shy2-2*-GR. **(D)**, Cross-sections (plastic embedding) of the uppermost part of Col0, *erl1*, *erl2* and *er* roots. 6-day-old plants were transferred for 2 days on Mock (left panels) or 1µM NAA (right panels) plates. **(E)**, Quantification of the number of secondary vascular (SV) tissues and xylem cells in (FigS3D). M indicates Mock and N indicates NAA. **(F)**, Cross-sections (plastic embedding) of the uppermost part of 12-day-old p*ER*::ER-YPet, *er erl1 erl2* (left panel), and p*PXY*::NSImb-vhhGFP4 in p*ER*::ER-YPet, *er erl1 erl2* (right panel) roots. The xylem vessels are highlighted in blue. **(G)**, Quantification of the number of secondary vascular tissues and xylem cells in (FigS3F). Black scale bars: 20µm. Count data was modelled with a generalized mixed model. Here the Poisson mix model was selected. P values were adjusted using Holm-Bonferroni correction.

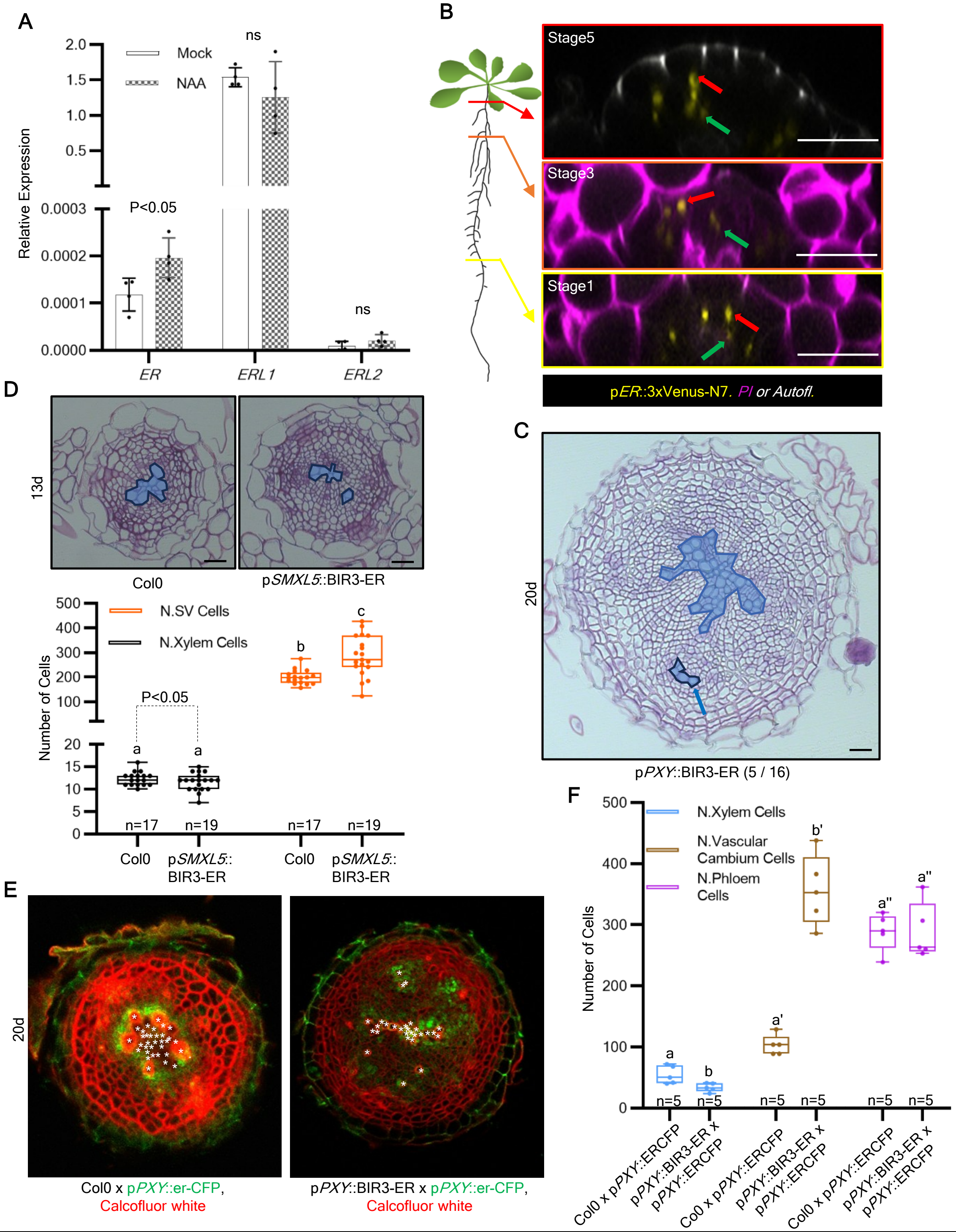


**Fig. S4. ER controls vascular cambium derivative formation. Related to Figure 2.**
**(A)**, Relative expression of *ER*, *ERL1* and *ERL2* in the Col0. 12-day-old plants were transferred for 1 days on Mock or 1µM NAA plates. **(B)**, Orthogonal view of z stacks of p*ER*::3xVenus-N7. 12-day-old roots at the positions corresponding to stage 1 (lower panel, before the initiation of periderm), stages 3 (middle panel, on the initiation of periderm), and stage 5 (up panel, after the initiation of periderm) of secondary growth. The red arrow indicates cork cambium / pericycle and the green arrow indicates vascular cambium. PI: Propidium Iodide. **(C)**, Cross-sections (plastic embedding) of the uppermost part of 20-day-old p*PXY*::BIR3-ER (right panel) roots. The blue arrow indicates the ectopic secondary xylem cells. The xylem vessels are highlighted in blue. **(D)**, Up-panels: cross-sections (plastic embedding) of the uppermost part of 13-day-old Col0 and Psmxl*5*::BIR3-ER. Lower panel: quantification of the number of secondary vascular (SV) tissues and xylem cells in (FigS4D). The xylem vessels are highlighted in blue. **(E)**, Vibratome-sections of the uppermost part of 20-day-old Col0 x p*PXY*::er-CFP (upper panel), and p*PXY*::BIR3-ER x p*PXY*::er-CFP (lower panel) F1 roots. White asterisk indicates xylem cells. **(F)**, Quantification of the number of xylem cells, vascular cambium cells and Phloem cells in (FigS4E). Black and white scale bars: 20µm. Count data was modelled with a generalized mixed model. Here the Poisson mix model was selected. P values were adjusted using Holm-Bonferroni correction. Count data was also analysed using the Student's T-test.

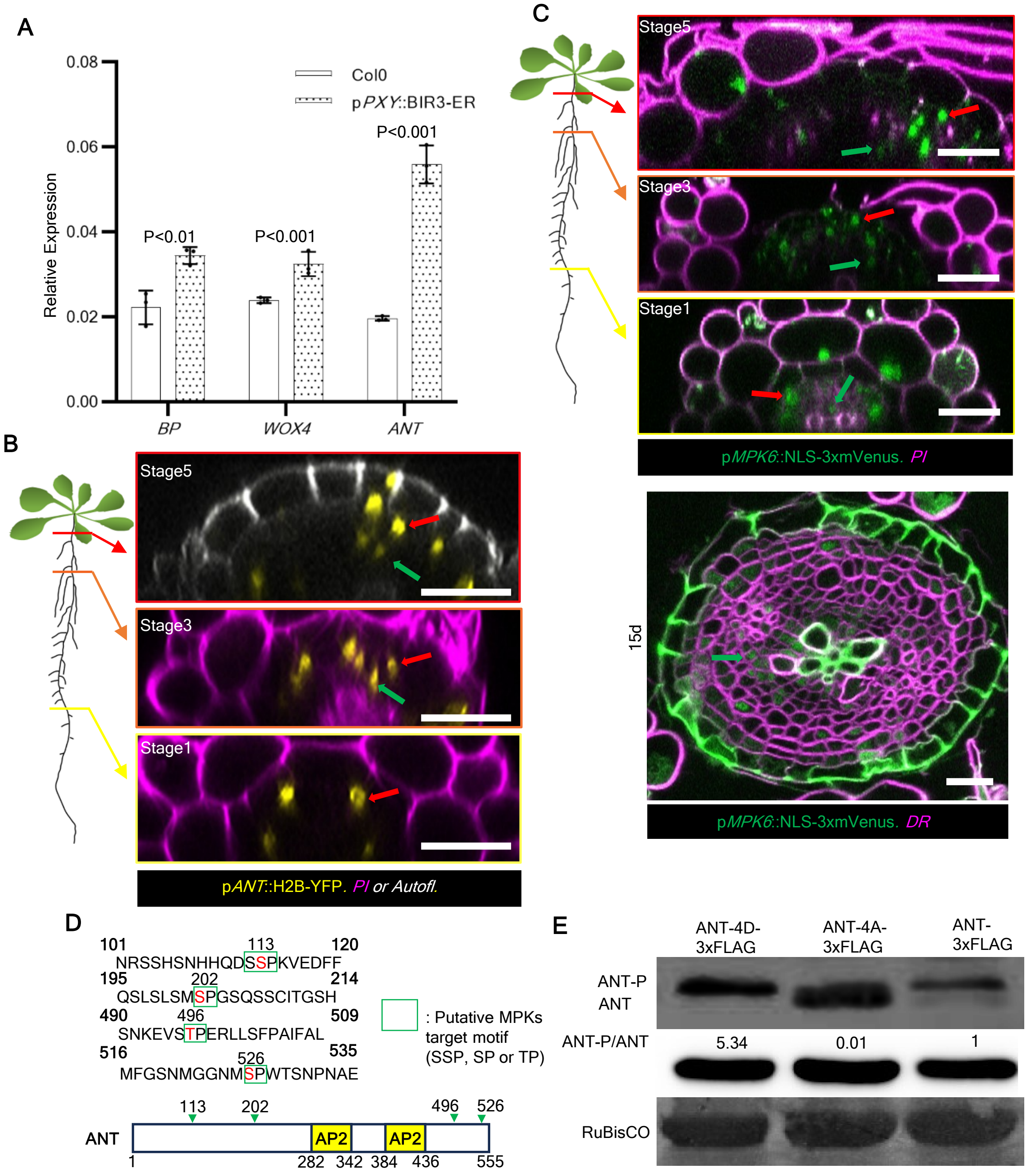


**Fig. S5. ANT is phosphorylated by MPK6. Related to Figure 2 and Figure3.**
**(A)**, Relative expression of *BP*, *WOX4*, and *ANT* in the Col0 and p*PXY*::BIR3-ER roots. 14-day-old roots for collection. **(B)**, Orthogonal view of Z-stacks of p*ANT*::H2B-YFP. 12-day-old roots at the positions corresponding to stage 1 (lower panel), stages 3 (middle panel), and stage 5 (up panel) of secondary growth. The red arrow indicates the cork cambium / pericycle and the green arrow indicates the vascular cambium. **(C)**, Upper panels: orthogonal view of Z-stacks of p*MPK6*::NLS-3xmVenus. 14-day-old roots at the positions corresponding to stage 1 (lower panel), stages 3 (middle panel), and stage 5 (up panel) of secondary growth. Lower panels: vibratome-sections of the uppermost part of 15-day-old p*MPK6*::NLS-3xmVenus. The red arrow indicates the cork cambium / pericycle and the green arrow indicates the vascular cambium. **(D)**, Putative MPK6 phosphorylation sites in the ANT sequence. ANT contains two AP2 domains. Four putative MPK6 phosphorylation residues (S113, S202, T496 and S526) were highlighted in the ANT protein. **(E)**, *In vivo* phosphorylation assays in *N. benthamiana*. The p*35S*::ANT-4D-3xFLAG (Phospho-mimic version of ANT), the p*35S*::ANT-4A-3xFLAG (Phospho-dead version of ANT) and the p*35S*::ANT-3xFLAG vectors were transiently expressed in leaves. Protein extracts were analysed on a Phos-tag/Normal SDS-PAGE gel and probed with an anti-Flag antibody. White scale bars: 20µm. PI: Propidium Iodide; DR: Direct Red. Count data was modelled with a generalized mixed model. Here the Poisson mix model or Poisson model was selected. P values were adjusted using Holm-Bonferroni correction.

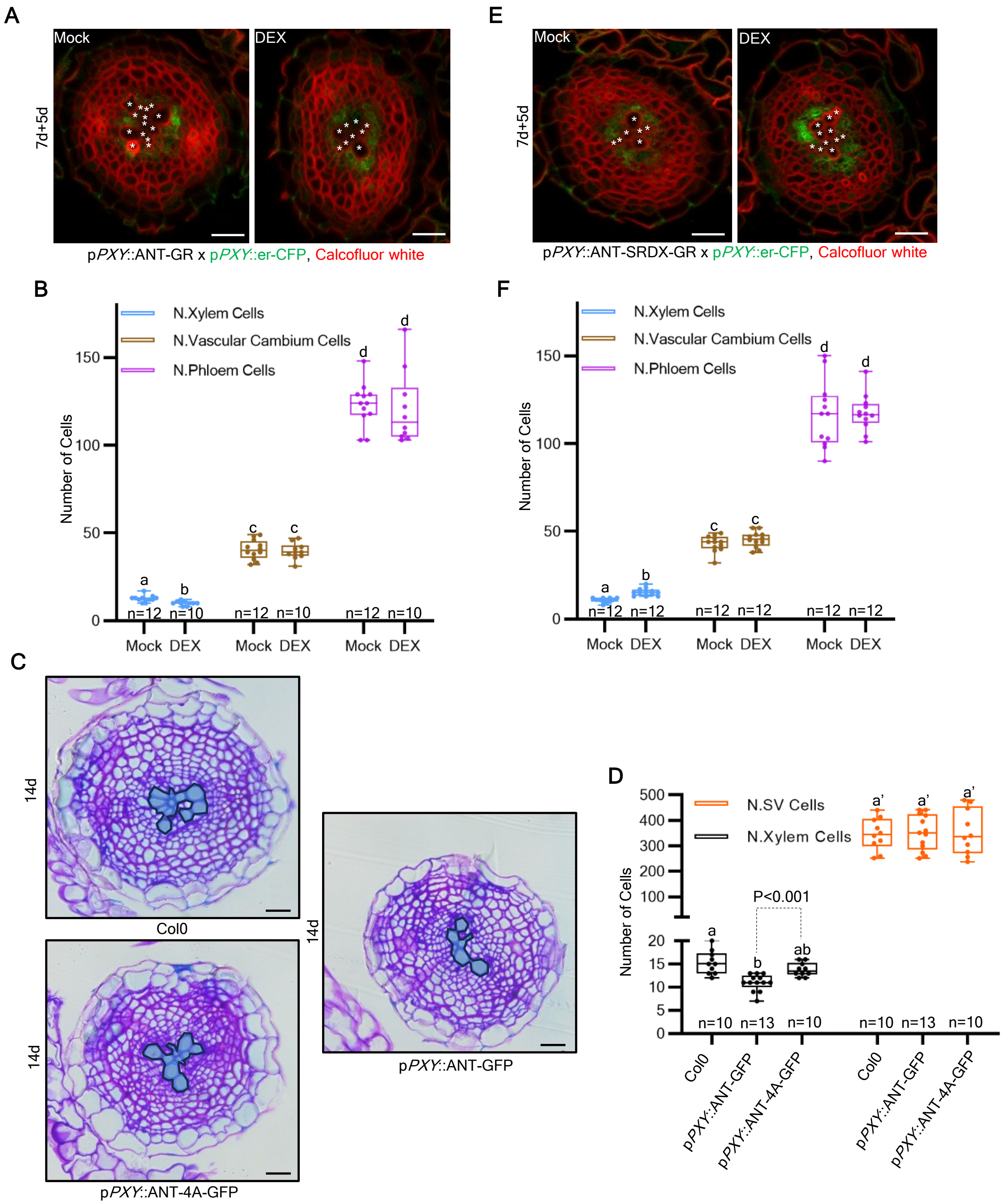


**Fig. S6. ANT acts downstream of ER signalling to control vascular cambium output. Related to Figure 4.**
**(A)**, Vibratome-sections of the uppermost part of p*PXY*::ANT-GR x p*PXY*::er-CFP F1 plants. 7-day-old plants were transferred to either Mock (left panel) or 10 µM DEX (right panel) plates for 5 days. White asterisk indicates xylem cells. **(B)**, Quantification of the number of xylem cells, vascular cambium cells and phloem cells in (FigS6A). **(C)**, Cross-sections (plastic embedding) of the uppermost part of 14-day-old Col0, and T1 of p*PXY*::ANT-GFP and p*PXY*::ANT-4A-GFP roots. The xylem vessels are highlighted in blue. **(D)**, Quantification of xylem cell number and secondary vascular (SV) tissues cell number in 14-day-old root shown in (FigS6C). **(E)**, Vibratome-sections of the uppermost part of p*PXY*::ANT-SRDX-GR x p*PXY*::er-CFP F1 plants. 7-day-old plants were transferred to either Mock (left panel) or 10 µM DEX (right panel) plates for 5 days. White asterisk indicates xylem cells. **(F)**, Quantification of the number of xylem cells, vascular cambium cells and phloem cells in (FigS6E). Black and white scale bars: 20µm. Count data was modelled with a generalized mixed model. Here the Poisson mix model or Poisson model was selected. P values were adjusted using Holm-Bonferroni correction. Count data was also analysed using the Student's T-test.

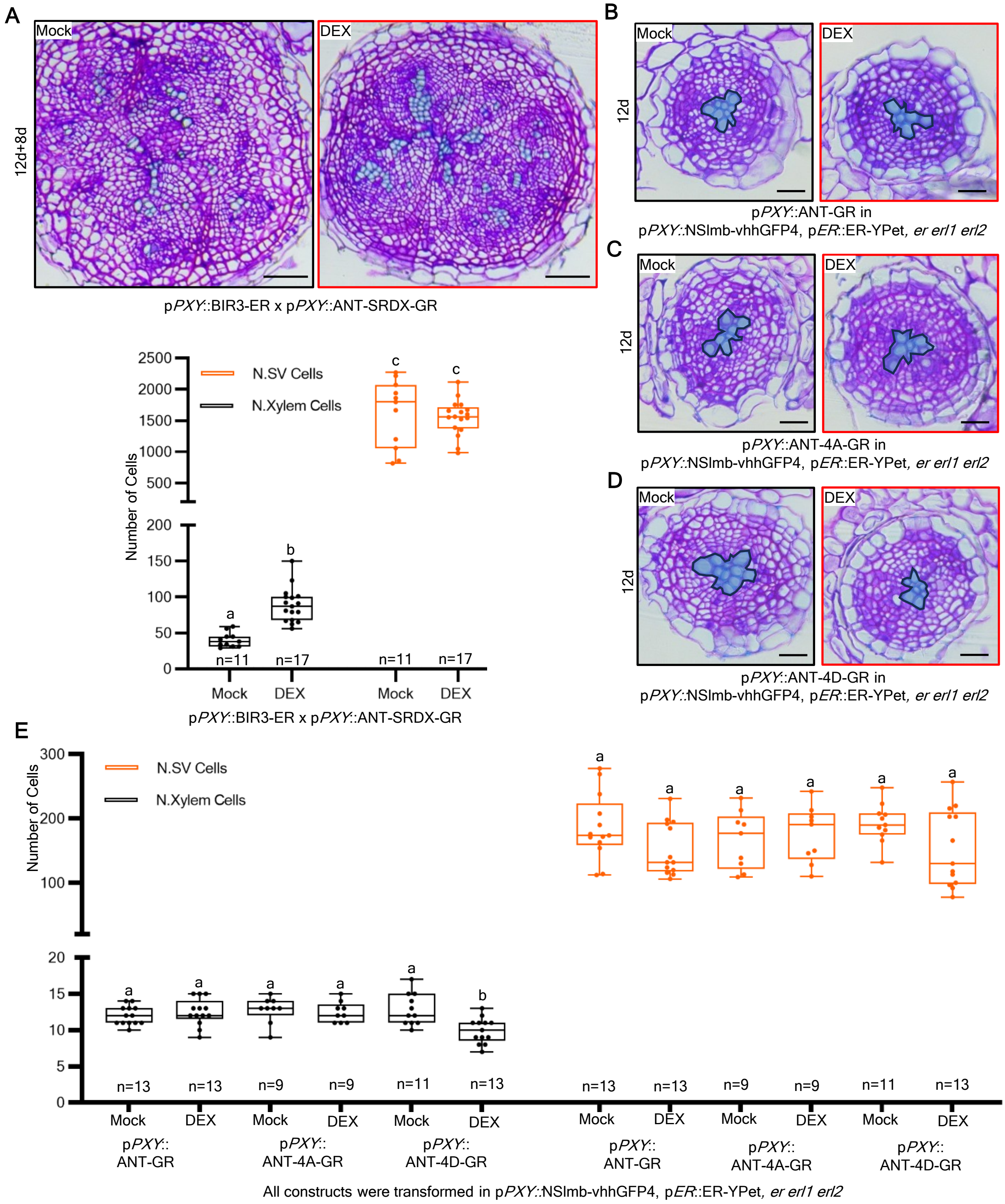


**Fig. S7. ANT phosphorylation depends on ER signalling. Related to Figure 4.**
**(A)**, Upper panels: root cross-sections of p*PXY*::BIR3-ER x p*PXY*::ANT-SRDX-GR. 12-day-old plants were transferred to either Mock (left panel) or 10 µM DEX (right panel) plates for 8 days. Lower panel: quantification of 20-day-old root in xylem cell number and secondary vascular tissues cell number of the experiment shown in (FigS7A). **(B-D)**, The root cross-sections of p*PXY*::ANT-GR (A), p*PXY*::ANT-4A-GR (B), and p*PXY*::ANT-4D-GR (C) in p*PXY*::NSlmb-vhhGFP4 in p*ER*::ER-YPet, *er erl1 erl2* plants. 12-day-old plants were grown on Mock (left panel) or 10µM DEX (right panel) plates. The xylem vessels are highlighted in blue. **(E)**, Quantification of 12-day-old root in xylem cell number and secondary vascular tissues cell number of the experiment shown in (FigS7B-D). Black scale bars: 20µm. Count data was modelled with a generalized mixed model. Here the Poisson mix model or Poisson model was selected. P values were adjusted using Holm-Bonferroni correction.

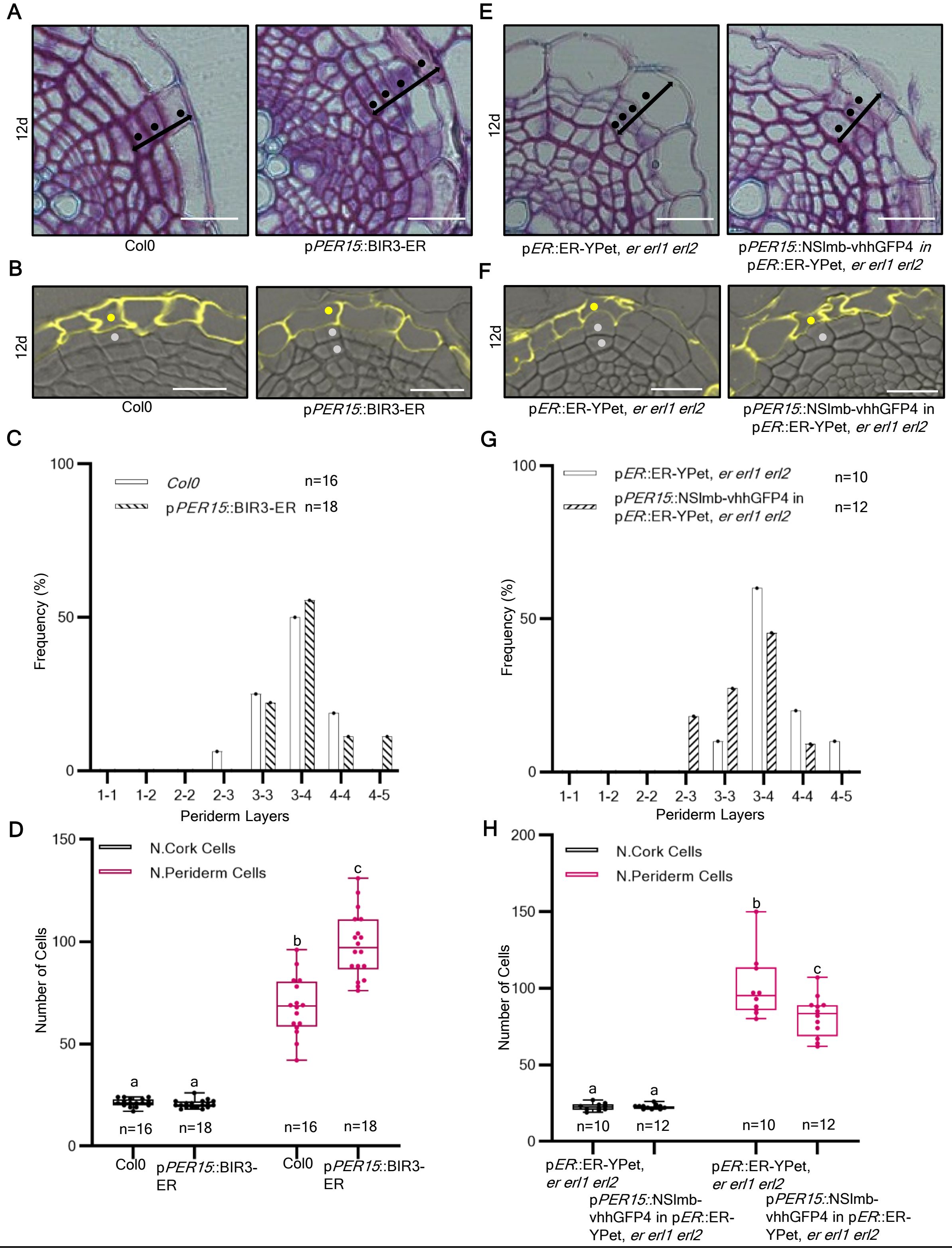


**Fig. S8. ER balances cork cambium output. Related to Figure 4.**
**(A)**, Cross-sections of the uppermost part of 12-day-old Col0 (left panel) and p*PER15*::BIR3-ER (right panel) roots. Black dots represent periderm cell layers. **(B)**, Cross-sections (plastic embedding) stained with Fluorol yellow (FY) of the uppermost part of 12-day-old Col0 (left panel) and p*PER15*::BIR3-ER (right panel) roots. Yellow dots represent suberized periderm layers and grey dots represent non-suberized periderm layers. **(C)**, Quantification of periderm layers number of the experiment shown in (Fig S7A). **(D)**, Quantification of the number of periderm and cork cells in (FigS7A). **(E)**, Cross-sections of the uppermost part of 12-day-old p*ER*::ER-YPet, *er erl1 erl2* (left panel), p*PER15*::NSlmb-vhhGFP4 in p*ER*::ER-YPet, *er erl1 erl2* (right panel) roots. Black dots represent periderm cell layers. **(F)**, Cross-sections (plastic embedding) stained with Fluorol yellow (FY) of the uppermost part of 12-day-old Col0 (left panel) and p*PER15*::BIR3-ER (right panel) roots. Yellow dots represent suberized periderm layers and grey dots represent non-suberized periderm layers. **(G)**, Quantification of periderm layers of the experiment shown in (Fig S7E). **(H)**, Quantification of the number of periderm and cork cells in (FigS7E). White scale bars: 20µm. Count data was modelled with a generalized mixed model. Here the Poisson mix model or Poisson model was selected. P values were adjusted using Holm-Bonferroni correction.

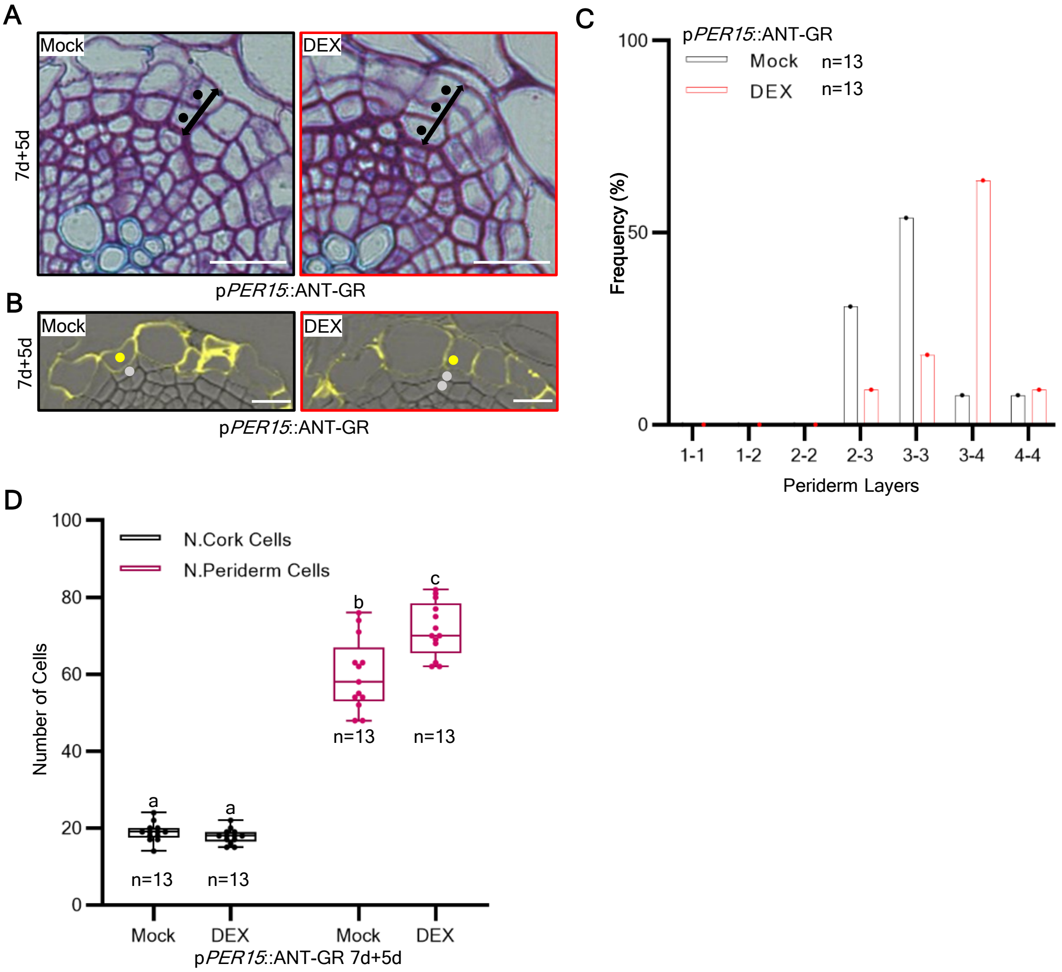


**Fig. S9. ANT balances cork cambium derivative formation. Related to Figure 4.**
**(A)**, Cross-sections of the uppermost part of 12-day-old p*PER15*::ANT-GR roots; 7-day-old plants were transferred for 5 days on Mock (left panel) or 10µM DEX (right panel) plates. Black dots represent periderm cell layers. **(B)**, Cross-sections (plastic embedding) stained with Fluorol yellow (FY) of the uppermost part of 12-day-old p*PER15*::ANT-GR roots. 7-day-old plants were transferred for 5 days on Mock (left panel) or 10µM DEX (right panel) plates. Yellow dots represent suberized periderm layers and grey dots represent non-suberized periderm layers. **(C)**, Quantification of periderm layers of the experiment shown in (FigS9A). **(D)**, Quantification of the number of periderm and cork cells in (FigS9A). White scale bars: 20µm. Count data was modelled with a generalized mixed model. Here the Poisson mix model was selected. P values were adjusted using Holm-Bonferroni correction.

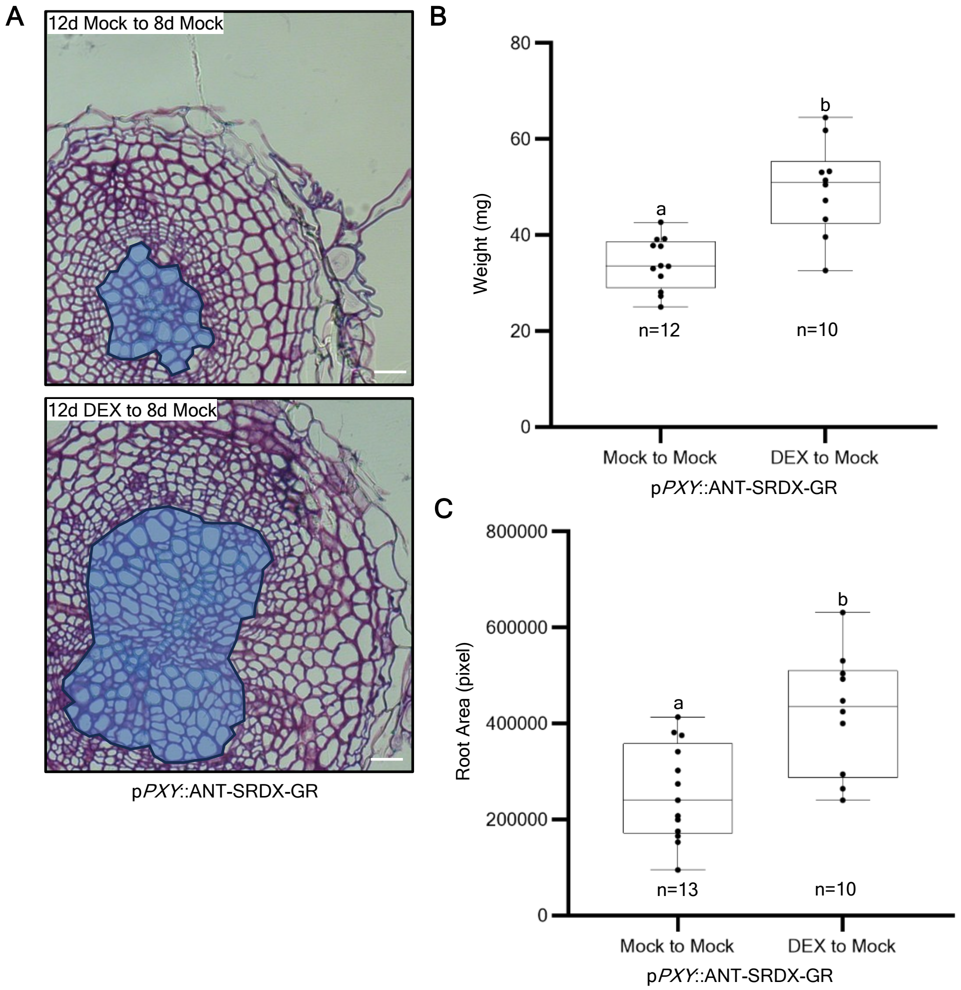


**Fig. S10. Extra secondary xylem cells enhance the plant growth. Related to Figure 4.**
**(A)**, The root cross-sections of p*PXY*::ANT-SRDX-GR plants grown in Mock / DEX plates for 20 day are shown. 12-day-old plants on Mock (up panel) or DEX (lower panel) plates and were transferred to Mock plates. The xylem vessels are highlighted in blue. **(B-C)**, Quantification of fresh weight of the shoot **(B)** and area of the root **(C)** is measured in the experiment shown in (**A**). White scale bars: 20μm. Count data was modelled with a generalized mixed model. Here the Poisson mix model was selected. P values were adjusted using Holm-Bonferroni correction.
